# Stress Granule-Related Genes during Embryogenesis of an Invertebrate Chordate

**DOI:** 10.1101/2024.02.26.582042

**Authors:** Laura Drago, Alessandro Pennati, Ute Rothbächer, Ryuji Ashita, Seika Hashimoto, Ryota Saito, Shigeki Fujiwara, Loriano Ballarin

## Abstract

Control of global protein synthesis through the assembly of stress granules (SGs) represents a strategy adopted by eukaryotic cells to face various stress conditions. TIAR, TTP and G3BP are key components of SGs, allowing the regulation of mRNA stability and thus controlling not only stress responses but also cell proliferation and differentiation. In the present work we aimed to investigate the role of *tiar*, *ttp* and *g3bp* during embryogenesis of the solitary ascidian *Ciona robusta*, both in physiological and stress conditions. We carried out CRISPR/Cas9 knockout to evaluate the effects on normal embryonic development, and gene reporter assays to study time and tissue specificity of gene expression, together with whole mount ISH and qRT-PCR. To induce acute stress conditions, we used iron and cadmium as “essential” and “non-essential” metals, respectively. Our results highlight, for the first time, the importance of *tiar, ttp* and *g3bp* in the control of development of mesendodermal tissue derivatives during embryogenesis of an invertebrate chordate.

**HIGHLIGHTS:**

- Mesenchyme, nervous cells, and endoderm are the embryonic regions mainly transcribing
*cr-tiarx1*, *cr-ttp* and *cr-g3bp2* during metal stress conditions.
- Only *cr-ttp* transcription is regulated at mid gastrula stage.
- The highest transcription levels are reached in hatching larvae after iron exposure.
- *cr-tiarx1*, *cr-ttp* and *cr-g3bp2* knockouts resulted in impaired embryogenesis.

## INTRODUCTION

To store energy for maintaining normal cellular homeostasis, eukaryotic cells have evolved sophisticated strategies, which implicate the attenuation of global protein synthesis through the formation of stress granules (SGs). The dynamic assembly of this membraneless foci in the cytoplasm, mediated by the overexpression of mRNA binding proteins, allows the temporary stop of translation initiation of selected mRNAs confined inside SGs, in order to prepare the cell for a future acute stress condition when mRNAs are rapidly released from SGs to codify the required proteins (Arribere et al. 2011; Liu and Qian, 2014; Protter and Parker, 2016; Somasekharan et al., 2020). SGs consist in a core structure of key mRNA binding proteins and mRNAs, connected each other by strong interactions, surrounded by a less concentrated and more dynamic shell, in which proteins and mRNAs can be easily exchanged with other type of cytoplasmic foci, such as processing bodies (p-bodies) (García-Mauriño et al., 2017), where mRNAs undergo degradation, germ cell granules (Mukherjee and Mukherjee, 2021), for maternal mRNA storage during early development, and neuronal granules (Söhnel and Brandt, 2023), important for synaptic remodeling. The regulation of mRNA stability, therefore, plays a key role not only in stress responses but also in cell proliferation, differentiation, and development.

TIA 1 related nucleolysin (TIAR) and Ras-GTPase-Activating Protein SH3-Domain-Binding Protein (G3BP) are components of SG core, whereas tristetraprolin (TTP) is a shell element. These proteins can recognize and bind AU-rich elements (AREs) at the 3′, functioning as deadenylation signal, or the 5’ untranslated regions (UTRs) of mRNAs for proteins involved in cell growth and differentiation, signal transduction, apoptosis, nutrient transport, and metabolism (e.g. *tnf-α, p38, stat5b, myc*) (Gallouzi et al., 1998; Piecyk et al., 2000; Dean et al., 2004; Vignudelli et al., 2010; Geng et al., 2015; Makita et al., 2021). The post-transcriptional control, operated by SGs, is allowed by the presence of RNA-Recognition Motifs (RRMs), in TIAR and G3BP, and CCCH tandem zinc finger (Znf) domains, in TTP (Cléry et al., 2008; von Roretz et al., 2011).

Mammalian TIAR has three RRMs, as documented: RRM1 confer the ability to interact with other RNA-binding proteins, RRM2 is the main DNA/RNA interaction motif, and RRM3 can regulate the TIAR export from nucleus to cytoplasm (Waris et al., 2014). A detailed description of TIAR can be found in our previous work (Drago et al., 2023). The importance of TIAR during embryogenesis was documented in mice (Sánchez-Jiménez and Izquierdo, 2013; Kharraz et al., 2010; Liu et al., 2023), where both the under- and the overexpression of the gene is associated with impaired embryo formation or lethality, and in *Caenorhabditis elegans* (Huelgas-Morales et al., 2016), in protection of the germline from heat stress. In addition, *tiar* knockout is known to cause infertility or impaired gametogenesis in adult mice (Beck et al., 1998; Piecyk et al., 2000).

G3BP is a site-specific ribonucleic acid endonuclease, able to bind the SH3 domain of Ras (apoptosis-inducing factor)-GTPase-activating protein (GAP) in serum-stimulated cells, as firstly reported by Parker et al. (1996). To exert its catalytic activity, G3BP requires phosphorylation sites. In proliferating cells, for example, G3BP is hypo-phosphorylated, leading to the loss of its ability to cleave AU-rich mRNA. In mammals, G3BP contains one carboxyl C-terminal RRM, with conserved hydrophobic amino acids in the ribonucleoprotein 1 (RNP1; eight amino acids) and ribonucleoprotein 2 (RNP2; six amino acids) sequences, both essential for the bond with RNA, one N-terminal nuclear transport factor 2 (NTF2) domain, associated with G3BP nuclear translocation and dimerization by mediating the Ran-GDP nuclear import through nucleoporins, and one N-terminal arginine-glycine-rich (RGG) region, which can be easily methylated allowing the regulation of SG disassembly. Conversely, the C-terminal region of G3BP can induce the phosphorylation of eucaryotic initiation factor 2 alpha (eIF2α), which reduces the availability of the eIF2–GTP–tRNAMet ternary complex necessary to initiate protein translation, so promoting the SG assembly (Kang et al., 2021; Tourrière et al., 2023).

Three homologous G3BP are known: G3BP1, G3BP2a and G3BP2b. G3BP1 differs from G3BP2 for a substitution (valine with isoleucine) at RNP2, and G3BP2a lacks 33 amino acids in its central proline-rich region respect to G3BP2b. Proline-rich motifs (PxxP) are required for the activation of protein kinase (PKR) important for the nucleation of SGs: G3BP1 has only one PxxP motif, whereas G3BP2a and G3BP2b have four and five PxxP motifs, respectively (Kang et al., 2021). G3BP2 contains more arginine residues in the RGG region than G3BP1, which possibly lead to different target specificities of mRNAs and make G3BP2 more prone to oligomerization. In mammals, G3BP1 is highly expressed in lung and kidney, whereas G3BP2 is highly expressed in the small intestine and brain (Kennedy at al., 2001). G3BP1 and G3BP2, through their NTF2 domain, can also form homo- or hetero-dimers, thus explaining their co-existence in SGs, since the dysfunction of one of them can negatively affect the other (Jin et al., 2022). The NTF2 domain plays an important role in viral replication, since it can bind to the viral motifs allowing the G3BP recruitment by the viral replication complex to evade the cellular immunity of the host (Götte et al., 2019; Wang and Merits, 2022). In mouse embryogenesis, the inactivation of G3BP leads to embryonic lethality or growth retardation and neonatal lethality (Zekri et al., 2005).

TTP, also known as TIS11 and ZFP36, promotes the poly(A) tail removal or deadenylation in AU-rich mRNAs, leading to their conservation or degradation, respectively in SGs and p-bodies, with partially or completely re-localization from p-bodies to SGs in stressed cells (Kedersha et al., 2005). The activation of *ttp* was first observed by Varnum et al. (1989) in murine Swiss 3T3 cells, as primary response to the tumor promoter tetradecanoyl phorbol acetate (TPA); TTP translocates from the nucleus to the cytoplasm upon stimulation with mitogen (Taylor et al., 1996). Two or more ZnF domains, able to bind Zn^2+^ with high affinity, were identified in TTP from mammals, *Xenopus*, *Drosophila,* and yeast: they possible allow the self-regulation of the protein by destabilizing its own mRNA. In addition, the Tis11D domain is involved in calcium signaling-induced apoptosis in B cells (Murata et al., 2005). In mammals, four homologous TTPs are known, the knockout of which cause systemic inflammatory syndrome in mice, due to the chronic increase of TNFα levels, and embryonic lethality. TTP negatively regulates hematopoietic/erythroid cell differentiation by promoting STAT5B mRNA decrease (Vignudelli et al., 2010). TTP is implicated in maternal mRNA turnover in mouse embryogenesis and the gene disruption leads to female infertility (Ramos et al., 2004) in mice, it is also important for yolk sac and placenta development (Stumpo et al., 2016). Through mRNA deadenylation or degradation, TTP plays a role in regulating immune functions and cell growth. It can act as tumor suppressor by inhibiting cell proliferation in breast cancer cells (Huang et al., 2016) and, through the modulation of the inflammatory-mediator *tnf-α* in macrophages, can regulate apoptotic events (Johnson and Blackwell, 2002; Holmes et al., 2012).

TIAR, G3BP and TTP are widely studied in vertebrates as molecular markers of SGs. In marine invertebrates the only information on molecular markers of SGs are found in our previous works, in which we investigated *tiar* and *ttp* expression during metal-induced stress conditions in the solitary ascidian *Ciona robusta* (Drago et al., 2021), and the role of TIAR during non-embryonic development of the colonial ascidian *Botryllus schlosseri*, characterized by cyclical generation changes (Drago et al., 2023). In the present work we aim to elucidate the functions of *tiar*, *ttp* and *g3bp* during the early development of *C. robusta*, both under physiological and stress conditions. We used clustered regularly interspaced short palindromic repeats (CRISPR)/Cas9 technique to study the effect on normal embryo development of *tiar*, *g3bp* and *ttp* knockouts, and gene reporter assay to study time and cellular specificity of gene expressions, reflecting the action of the regulatory sequences, in embryos under control and stress conditions due to exposure to iron and cadmium, as “essential” and “non-essential” metals, respectively. *tiar*, *g3bp* and *ttp* expressions were analyzed through whole mount *in situ* hybridization (ISH) and quantitative real-time PCR (qRT-PCR).

## RESULTS

### GENE AND PROTEIN ORGANIZATION IS HIGHLY CONSERVED ACROSS METAZOANS

Gene and protein organizations of C*. robusta* TIAR and TTP (Cr*-*TIARx1 and Cr-TTP, respectively) were described in a previous paper (Drago et al., 2021). The Cr-g3bp_PCRFw and Cr-g3bp_PCRRv primers allowed us to amplify a region of 853 nt, whose sequencing confirmed the putative *g3bp* sequence that we found in ANISEED database (transcript ID: KY2019:KY.Chr1.2038.v1.SL1-1). This region, covering most of the coding DNA sequence (CDS) (Figure S1), showed, in NCBI database, the highest similarity (100%) and identity (97.62%) with *Ciona intestinalis* G3BP2 (GeneBank accession number: XM_002126499.5). The *in-silico* reconstruction resulted in a whole transcript of 1797 nt, we named *cr-g3bp2*, with 5′-UTR and 3′-UTR regions of 72 and 360 nt, respectively, and the CDS of 1365 nt, encoding a putative protein of 454 amino acids with a deduced molecular weight of 51.6 kDa (Figure S1). The gene is located in chromosome 1, as revealed by Ghost database, and organized in nine exons (length in nt from exon one to nine: 98, 82, 174, 103, 188, 174, 232, 113, 201) and eight introns (length in nt from intron one to eight: 368, 226, 1700, 463, 418, 211, 308, 229).

The multi-alignment analysis on G3BP2 amino acid sequences of tunicates (*Phallusia mammillata*, *Ciona savignyi*, *C. robusta*, *Halocynthia roretzi*, *Botrylloides leachii*), *Branchiostoma floridae, Latimeria chalumnae* and *Patella vulgata*, the last three as representative of cephalochordates, vertebrates and spiralian invertebrates respectively, shows: i) the conserved N-terminal NTF2 domain (50.4% of identical amino acids among sequences), ii) the C-terminal RRM domain (33.8% of identical amino acids among sequences), iii), the central acid-proline-rich region and iv) the C-terminal arginine- and glycine-rich region (Figure S2). In the RNP2 of RRM domain, the valine (V) residue was present: it allows to classify all the analyzed amino acid sequences, including that of *C. robusta,* as G3BP2. In the sequence of *Branchiostoma floridae*, which is the only G3BP sequence from cephalochordates found in NCBI, V is substituted by isoleucine (I), which is a prerogative of G3BP1. Despite this, among the examined non-tunicate species, Cr-G3BP2 showed the highest identity (47.4%) and similarity (68%) with that of *Branchiostoma floridae* (E-value 1.8e-40). Among all the species, the Cr-G3BP2 amino acid sequence showed the highest identity (78.4%) and similarity (90.1%) with that of *Ciona savignyi* (E-value 2.9e-107), as revealed by LALIGN tool (Table S2).

### GENE REPORTER ASSAYS SHOW METAL INDUCED mRNA ACCUMULATION IN MESENDODERM AND CENTRAL NERVOUS SYSTEM

The regulation of *tiarx1*, *ttp* and *g3bp2* transcription was analyzed in mid gastrula, mid tailbud II and hatching larva embryos of *C. robusta*. At mid gastrula stage, only the promoter of *cr*-*ttp* resulted active in the regions of neural plate and muscle precursors, with a higher involvement of neural plate and muscle cells in metal treated embryos (Figure 1).

**Figure 1.**
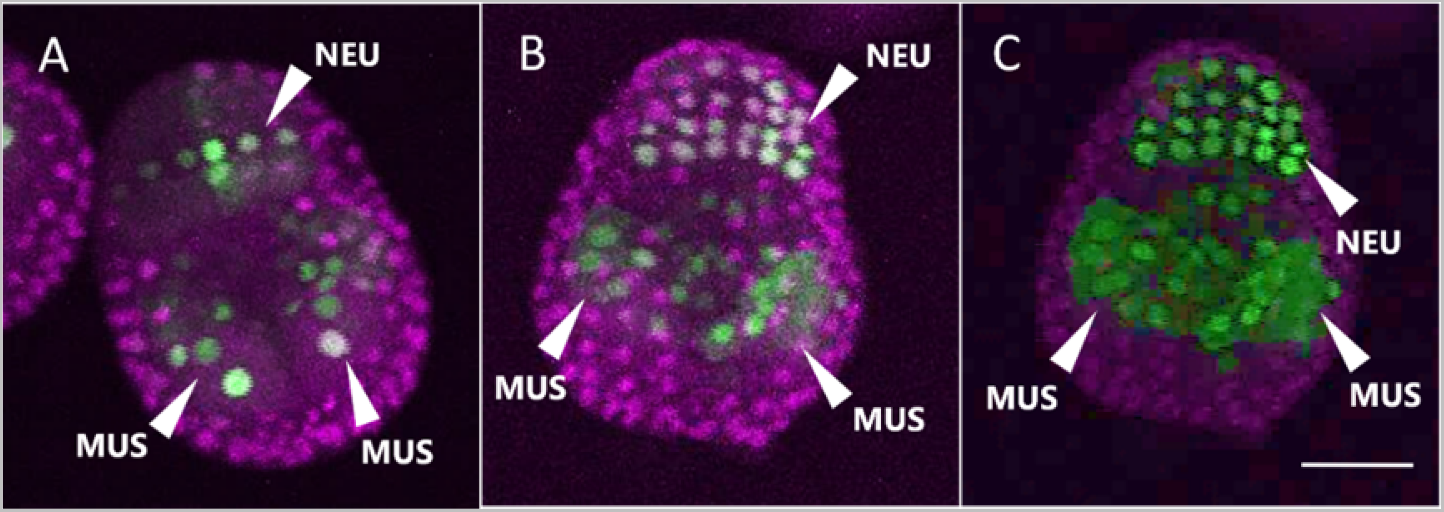
Gene reporter assay in mid gastrula embryos, after electroporation of pTtp>LacZ (A, B, C) (green), together with Fog>H2B:mCherry plasmid marking successful uptake of the electroporation mix (purple). Embryos are reported in control conditions (A), after cadmium exposure (B) and after iron exposure (C). MUS: muscle cells; NEU: neural plate. Scale bar: 50 μm.

In mid tailbud II stage, in all the experimental conditions (Ctr, Cd, Fe), the promoters were active in the mesenchyme. The activation was visible, for *cr-g3bp2* and *cr-ttp*, also in the nerve cord and other structures of the tail when embryos were exposed to Fe. *cr-g3bp2* expanded its activation area to the presumptive endoderm, brain, nerve cord and in other structures of the tail when embryos were exposed to Cd. As for *cr-tiar*, in addition to the mesenchyme, it resulted active also in the region of the brain after the exposure to Cd, and in the endodermal region after the exposure to Fe (Figure 2 and Table 1, 2, 3).

**Figure 2.**
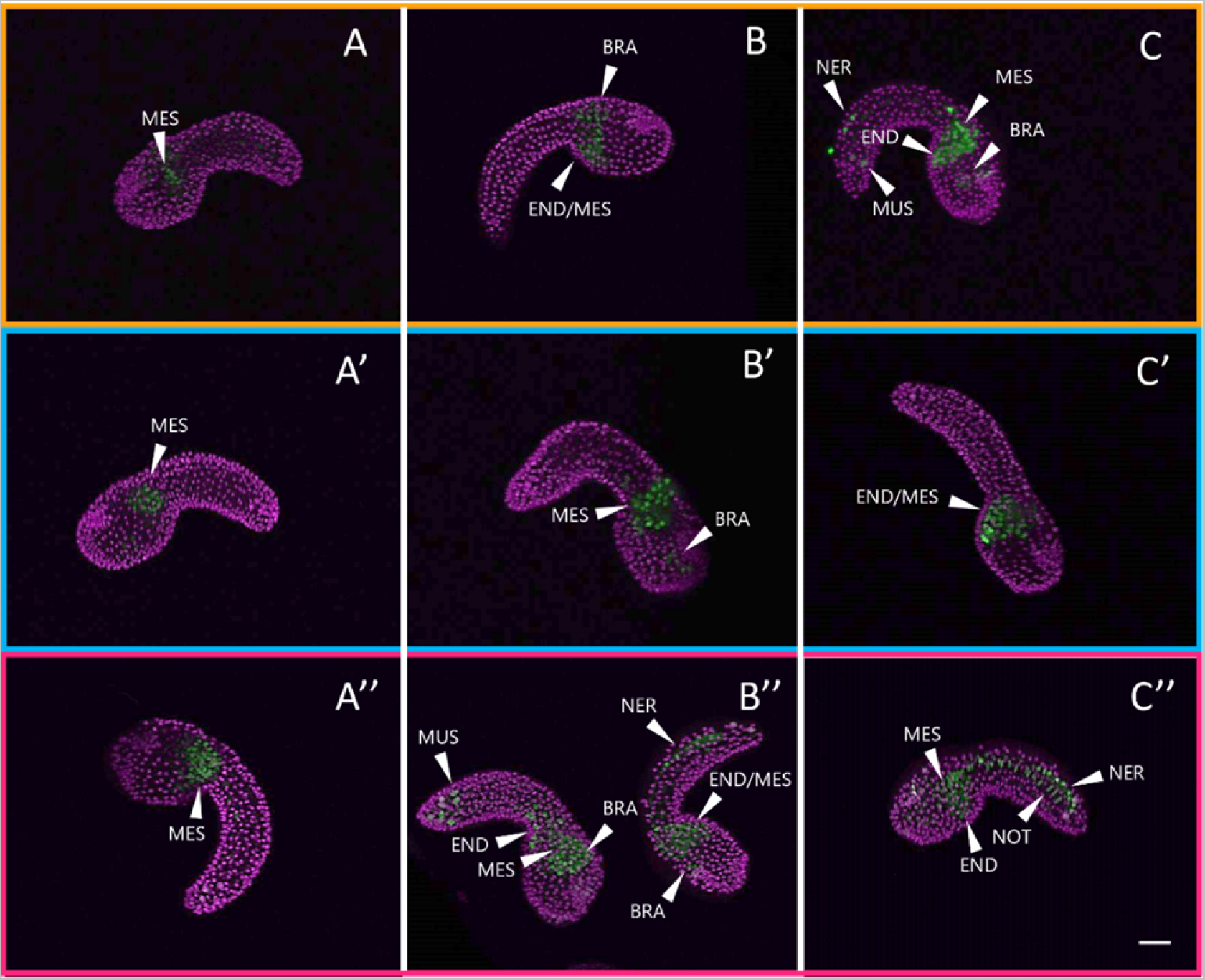
Gene reporter assay in mid tailbud II embryos, after electroporation of pG3bp>LacZ (A, B, C in orange box), pTiar>LacZ (A’, B’, C’ in blue box), or pTtp>LacZ (A’’, B’’, C’’ in red box) (green), together with Fog>H2B:mCherry plasmid marking successful uptake of the electroporation mix (purple). Embryos are reported in control conditions (A, A’, A’’), after cadmium exposure (B, B’; B’’) and after iron exposure (C, C’, C’’). MES: mesenchyme; END: endoderm; BRA: brain; NER: nerve cord; NOT: notochord; MUS: muscle cells. Scale bar: 100 μm.

**Table 1.** Mid tailbud II embryo counts according to marked tissue by gene reporter assays as regards *cr*-*g3bp2* expression regulation, after metal (Cd, Fe) and non-metal exposures: mesenchyme, endoderm (endoderm in the head together with endodermal strand in the tail), nervous system (brain in the head together with nerve cord in the tail), notochord and muscle cells. Χ^2^ was considered to compare frequencies of metal-treated and control embryos. Fog>H2B:mCherry refer to embryo counts without the examined tissue marked. In bold, percentage of embryos in which a statistically significant increase in tissue marked is present respect to control. ns: not significant.

| Readout | Ctr |  | Cd |  |  | Fe |  |  |
| --- | --- | --- | --- | --- | --- | --- | --- | --- |
|  | Embryo count | % with marked tissue | Embryo count | % with marked tissue | Chi squared (Ctr vs Cd) | Embryo count | % with marked tissue | Chi squared (Ctr vs Fe) |
| *Fog>H2B:mCherry* | 0 | 100% | 0 | 100% | ns | 0 | 100% | ns |
| *G3bp>LacZ* mesenchyme | 38 |  | 41 |  |  | 40 |  |  |
| *Fog>H2B:mCherry* | 27 | 29% | 29 | 29% | 0,6249 | 30 | 25% | 0,5148 |
| *G3bp>LacZ* endoderm | 11 |  | 12 |  |  | 10 |  |  |
| *Fog>H2B:mCherry* | 24 | 37% | 27 | 34% | 0,5403 | 17 | **58%** | **0,005146** |
| ***G3bp>LacZ* nervous system** | 14 |  | 14 |  |  | 23 |  |  |
| *Fog>H2B:mCherry* | 36 | 5% | 40 | 2% | 0,3311 | 40 | 0% | 0,1179 |
| *G3bp>LacZ* notocord | 2 |  | 1 |  |  | 0 |  |  |
| *Fog>H2B:mCherry* | 31 | 18% | 36 | 12% | 0,2405 | 31 | 23% | 0,4497 |
| *G3bp>LacZ* muscle cells | 7 |  | 5 |  |  | 9 |  |  |

**Table 2.** Mid tailbud II embryo counts according to marked tissue by gene reporter assays as regards *cr*-*tiarx1* expression regulation, after metal (Cd, Fe) and non-metal exposures: mesenchyme, endoderm (endoderm in the head together with endodermal strand in the tail), nervous system (brain in the head together with nerve cord in the tail), notochord and muscle cells. Χ^2^ was considered to compare frequencies of metal-treated and control embryos. Fog>H2B:mCherry refer to embryo counts without the examined tissue marked. In bold, percentage of embryos in which a statistically significant increase in tissue marked is present respect to control. ns: not significant.

| Readout | Ctr |  | Cd |  |  | Fe |  |  |
| --- | --- | --- | --- | --- | --- | --- | --- | --- |
|  | Embryo count | % with marked tissue | Embryo count | % with marked tissue | Chi squared (Ctr vs Cd) | Embryo count | % with marked tissue | Chi squared (Ctr vs Fe) |
| *Fog>H2B:mCherry* | 0 | 100% | 0 | 100% | ns | 0 | 100% | ns |
| *Tiar>LacZ* mesenchyme | 42 |  | 23 |  |  | 47 |  |  |
| *Fog>H2B:mCherry* | 34 | 19,05% | 19 | 17% | 0,003330 | 38 | 19,15% | 0,4402 |
| *Tiar>LacZ* endoderm | 8 |  | 4 |  |  | 9 |  |  |
| *Fog>H2B:mCherry* | 33 | 21,43% | 10 | 57% | 2,443E-05 | 29 | 38,30% | 0,002072 |
| ***Tiar>LacZ* nervous system** | 9 |  | 13 |  |  | 18 |  |  |
| *Fog>H2B:mCherry* | 41 | 2,38% | 22 | 4% | 0,003004 | 46 | 2,13% | 0,4349 |
| *Tiar>LacZ* nothocord | 1 |  | 1 |  |  | 1 |  |  |
| *Fog>H2B:mCherry* | 38 | 9,52% | 22 | 4% | 0,002719 | 45 | 4,26% | 0,1302 |
| *Tiar>LacZ* muscle cells | 4 |  | 1 |  |  | 2 |  |  |

**Table 3.** Mid tailbud II embryo counts according to marked tissue by gene reporter assays as regards *cr*-*ttp* expression regulation, after metal (Cd, Fe) and non-metal exposures: mesenchyme, endoderm (endoderm in the head together with endodermal strand in the tail), nervous system (brain in the head together with nerve cord in the tail), notochord and muscle cells. Χ^2^ was considered to compare frequencies of metal-treated and control embryos. Fog>H2B:mCherry refer to embryo counts without the examined tissue marked. In bold, percentage of embryos in which a statistically significant increase in tissue marked is present respect to control. ns: not significant.

| Readout | Ctr |  | Cd |  |  | Fe |  |  |
| --- | --- | --- | --- | --- | --- | --- | --- | --- |
|  | Embryo count | % with marked tissue | Embryo count | % with marked tissue | Chi squared (Ctr vs Cd) | Embryo count | % with marked tissue | Chi squared (Ctr vs Fe) |
| *Fog>H2B:mCherry* | 0 | 100,00% | 0 | 100,00% | ns | 0 | 100% | ns |
| *Ttp>LacZ* mesenchyme | 43 |  | 37 |  |  | 14 |  |  |
| *Fog>H2B:mCherry* | 29 | 32,60% | 28 | 24,32% | 0,1773 | 9 | **36%** | **9,653E-06** |
| ***Ttp>LacZ* endoderm** | 14 |  | 9 |  |  | 5 |  |  |
| *Fog>H2B:mCherry* | 20 | 53,49% | 17 | 54,05% | 0,3590 | 6 | **57,14%** | **9,634E-06** |
| ***Ttp>LacZ* nervous system** | 23 |  | 20 |  |  | 8 |  |  |
| *Fog>H2B:mCherry* | 28 | 11,63% | 35 | 5,41% | 0,05954 | 13 | 7,14% | 0,0008024 |
| *Ttp>LacZ* nothocord | 5 |  | 2 |  |  | 1 |  |  |
| *Fog>H2B:mCherry* | 32 | 25,58% | 17 | **54,05%** | **0,000148** | 7 | **50,00%** | **4,627E-06** |
| ***Ttp>LacZ* muscle cells** | 11 |  | 20 |  |  | 7 |  |  |

In hatching larvae, the tail did not show any sign of activity of the considered genes. Conversely, in the trunk, all the three genes were active in the endodermal region; *cr-g3bp2* and *cr-tiar* were active also in the mesenchyme. In Cd-treated specimens, the activation area increased in size in the case of *cr-tiar* and spread to the brain in the case of *cr-g3bp2*; no effects were visible for *cr-ttp*. When exposed to Fe, hatching larvae showed a general increase of the activation area for all the considered genes which spread in the trunk, and in the case of *cr-g3bp* also included the tail nerve cord (Figure 3 and Table 4, 5, 6).

**Figure 3.**
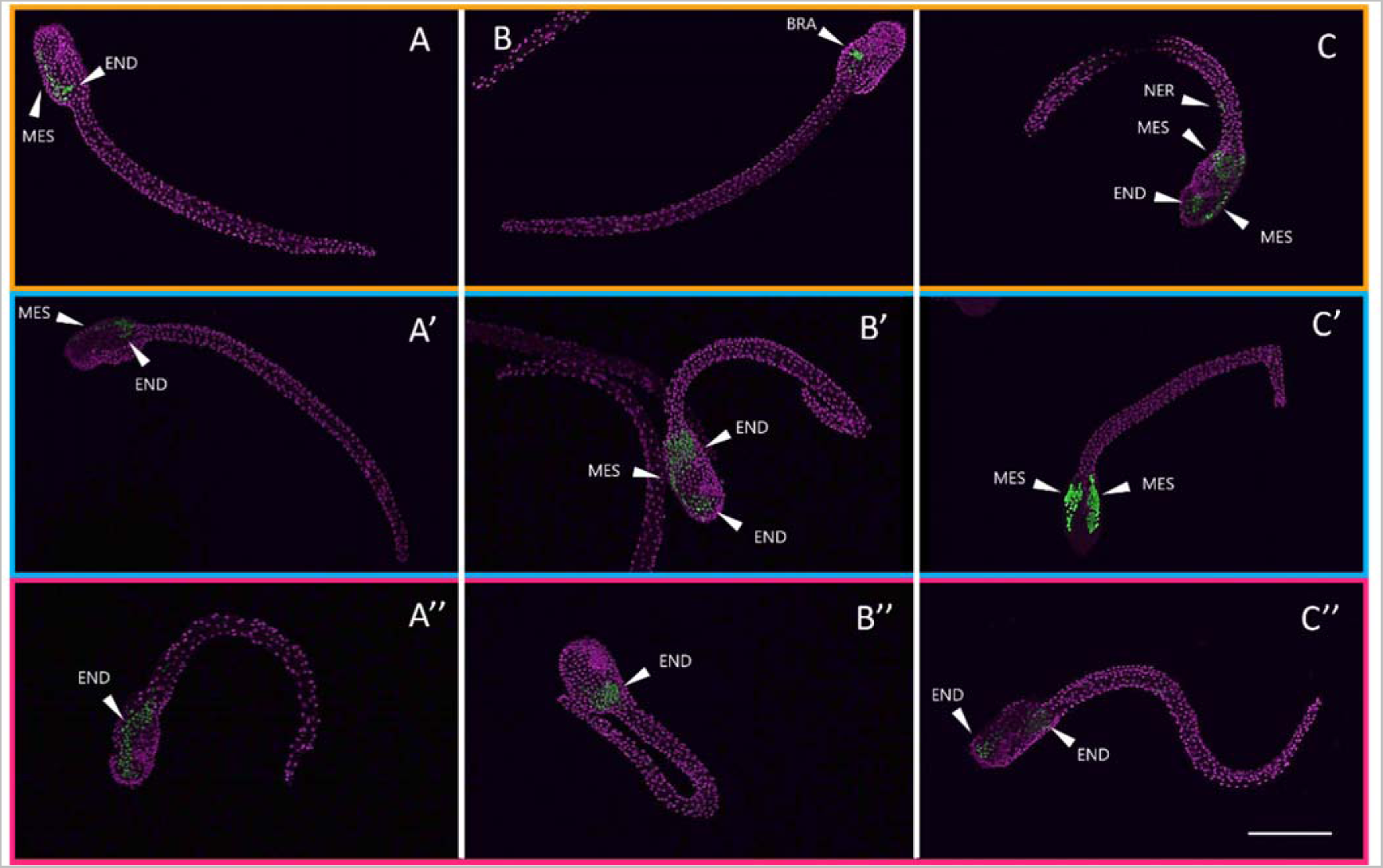
Gene reporter assay in hatching larva, after electroporation of pG3bp>LacZ (A, B, C, in orange box), pTiar>LacZ (A’, B’, C’, in blue box) or pTtp>LacZ (A’’, B’’, C’’, in red box) (green), together with Fog>H2B:mCherry plasmid marking successful uptake of the electroporation mix (purple). Embryos are reported in control conditions (A, A’, A’’), after cadmium exposure (B, B’; B’’) and after iron exposure (C, C’, C’’). MES: mesenchyme; END: endoderm; BRA: brain; NER: nerve cord. Scale bar: 100 μm.

**Table 4.** Hatching larva embryo counts according to marked tissue by gene reporter assays as regards *cr*-*g3bp2* expression regulation, after metal (Cd, Fe) and non-metal exposures: mesenchyme, endoderm (endoderm in the head together with nerve cord in the tail) and nervous system (brain in the head). Χ^2^ was considered to compare frequencies of metal-treated and control embryos. Fog>H2B:mCherry refer to embryo counts without the examined tissue marked. In bold, percentage of embryos in which a statistically significant increase in tissue marked is present respect to control.

| Readout | Ctr |  | Cd |  |  | Fe |  |  |
| --- | --- | --- | --- | --- | --- | --- | --- | --- |
|  | Embryo count | % with marked tissue | Embryo count | % with marked tissue | Chi squared (Ctr vs Cd) | Embryo count | % with marked tissue | Chi squared (Ctr vs Fe) |
| *Fog>H2B:mCherry* | 28 | 39,13% | 20 | 47,37% | 0,1306 | 54 | 37,93% | 1,419E-09 |
| *G3bp>LacZ* mesenchyme | 18 |  | 18 |  |  | 33 |  |  |
| *Fog>H2B:mCherry* | 32 | 30,43% | 22 | 42,11% | 0,06478 | 50 | **42,53%** | **4,461E-12** |
| ***G3bp>LacZ* endoderm** | 14 |  | 16 |  |  | 37 |  |  |
| *Fog>H2B:mCherry* | 32 | 30,43% | 34 | 10,53% | 0,007020 | 70 | 19,54% | 1,331E-11 |
| *G3bp>LacZ* nervous system | 14 |  | 4 |  |  | 17 |  |  |

**Table 5.** Hatching larva embryo counts according to marked tissue by gene reporter assays as regards *cr*-*tiarx1* expression regulation, after metal (Cd, Fe) and non-metal exposures: mesenchyme, endoderm (endoderm in the head together with nerve cord in the tail) and nervous system (brain in the head). Χ^2^ was considered to compare frequencies of metal-treated and control embryos. Fog>H2B:mCherry refer to embryo counts without the examined tissue marked. In bold, percentage of embryos in which a statistically significant increase in tissue marked is present respect to control.

| Readout | Ctr |  | Cd |  |  | Fe |  |  |
| --- | --- | --- | --- | --- | --- | --- | --- | --- |
|  | Embryo count | % with marked tissue | Embryo count | % with marked tissue | Chi squared (Ctr vs Cd) | Embryo count | % with marked tissue | Chi squared (Ctr vs Fe) |
| *Fog>H2B:mCherry* | 52 | 33,33% | 31 | **44,64%** | **0,004** | 25 | **48,98%** | **0,000167** |
| ***Tiar>LacZ* mesenchyme** | 26 |  | 25 |  |  | 24 |  |  |
| *Fog>H2B:mCherry* | 48 | 38,46% | 34 | 39,28% | 0,013 | 30 | 38,78% | 0,01024 |
| *Tiar>LacZ* endoderm | 30 |  | 22 |  |  | 19 |  |  |
| *Fog>H2B:mCherry* | 56 | 28,21% | 47 | 16,07% | 0,003 | 43 | 12,24% | 0,000129 |
| *Tiar>LacZ* nervous system | 22 |  | 9 |  |  | 6 |  |  |

**Table 6.** Hatching larva embryo counts according to marked tissue by gene reporter assays as regards *cr*-*ttp* expression regulation, after metal (Cd, Fe) and non-metal exposures: mesenchyme, endoderm (endoderm in the head together with nerve cord in the tail) and nervous system (brain in the head). Χ^2^ was considered to compare frequencies of metal-treated and control embryos. Fog>H2B:mCherry refer to embryo counts without the examined tissue marked. In bold, percentage of embryos in which a statistically significant increase in tissue marked is present respect to control.

|  | Ctr |  | Cd |  |  | Fe |  |  |
| --- | --- | --- | --- | --- | --- | --- | --- | --- |
| Readout | Embryo count | % with marked tissue | Embryo count | % with marked tissue | Chi squared (Ctr vs Cd) | Embryo count | % with marked tissue | Chi squared (Ctr vs Fe) |
| *Fog>H2B:mCherry* | 12 | 31,91% | 23 | **39,47%** | **0,001496** | 8 | **50,00%** | **0,03197** |
| ***Ttp>LacZ* mesenchyme** | 15 |  | 15 |  |  | 8 |  |  |
| *Fog>H2B:mCherry* | 38 | 19,15% | 20 | **47,37%** | **2,833E-05** | 10 | **37,50%** | **3,304E-06** |
| ***Ttp>LacZ* endoderm** | 9 |  | 18 |  |  | 6 |  |  |
| *Fog>H2B:mCherry* | 33 | 29,79% | 28 | 26,32% | 0,1680 | 14 | 12,50% | 4,083E-06 |
| *Ttp>LacZ* nervous system | 14 |  | 10 |  |  | 2 |  |  |

### WHOLE MOUNT ISH CONFIRMS ECTOPIC TRANSCRIPT ENRICHMENT BY METAL STRESSORS

The location of *cr-tiarx1*, *cr-ttp* and *cr-g3bp2* transcription was verified at mid gastrula and mid tailbud II stage, in which the tunic is not formed yet: in this way we were able to avoid the background signal. At the mid gastrula stage, the only positive signal was observed for *cr-ttp*, consistent with the early activation of the reporter gene. The signal was detected specifically in the neural plate for all the experimental conditions (Figure 4). *cr-tiarx1* and *cr-g3bp2* were not expressed at the mid gastrula stage (Figure 5 A and E). Conversely, all the studied genes were expressed at mid tailbud II stage, in mesenchyme and nervous system, also involving the nerve cord for *g3bp2* and *tiarx1*, after Cd and Fe exposure, respectively (Figure 4 and 5). In the case of *ttp*, also the endoderm of the head was involved in its expression at mid tailbud II stage (Figure 4). The absence of expression after the use of sense riboprobes was previously tested (Drago et al., 2021).

**Figure 4.**
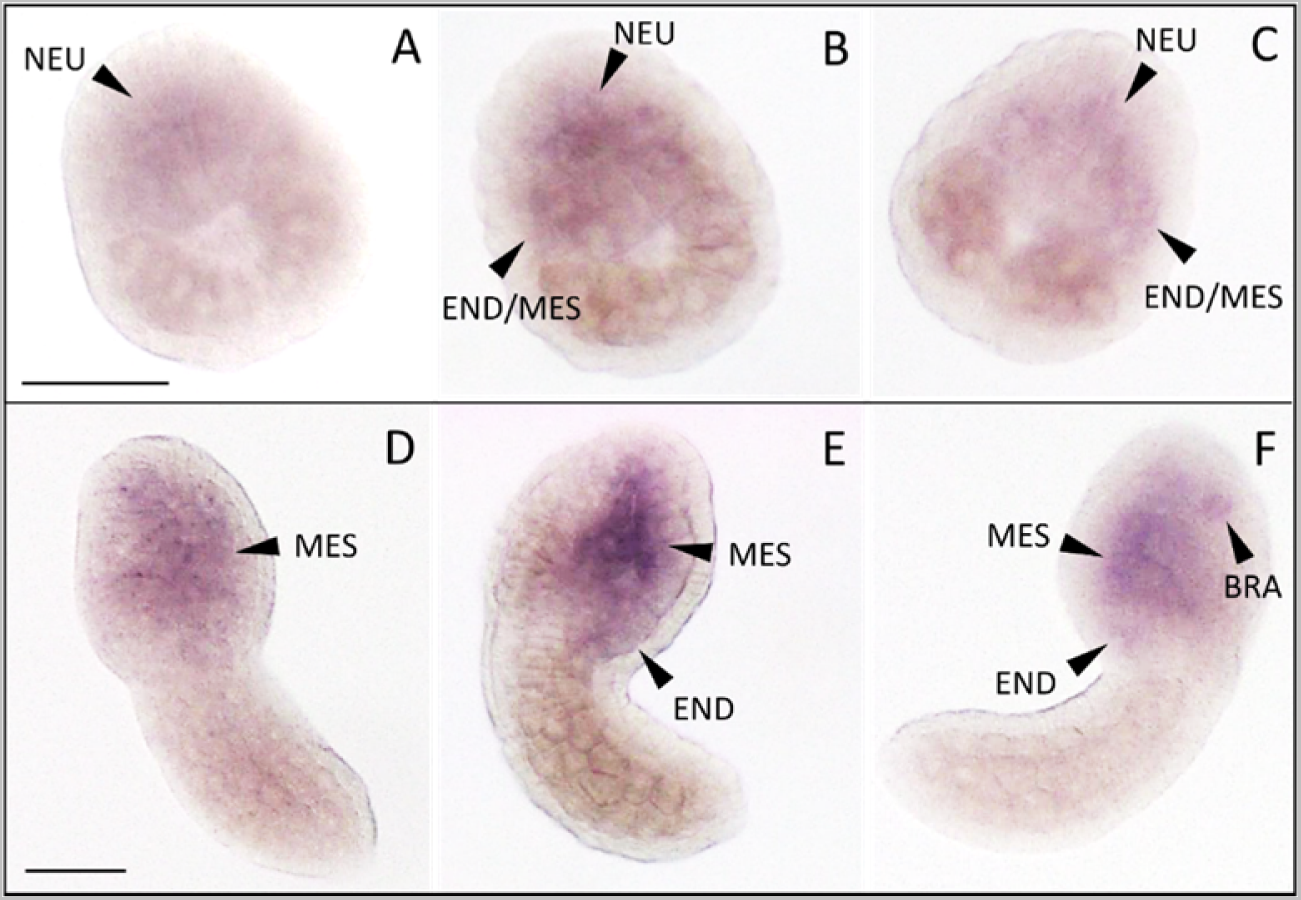
Whole mount ISH on mid gastrula (A-C) and mid tailbud II (D-F) embryos, with *cr-ttp* riboprobe, in control condition (A, D) or in presence of Fe (B, E) or Cd (C, F). NEU: neural plate; END: endoderm; MES: mesenchyme. Scale bar: 100 µm.

**Figure 5.**
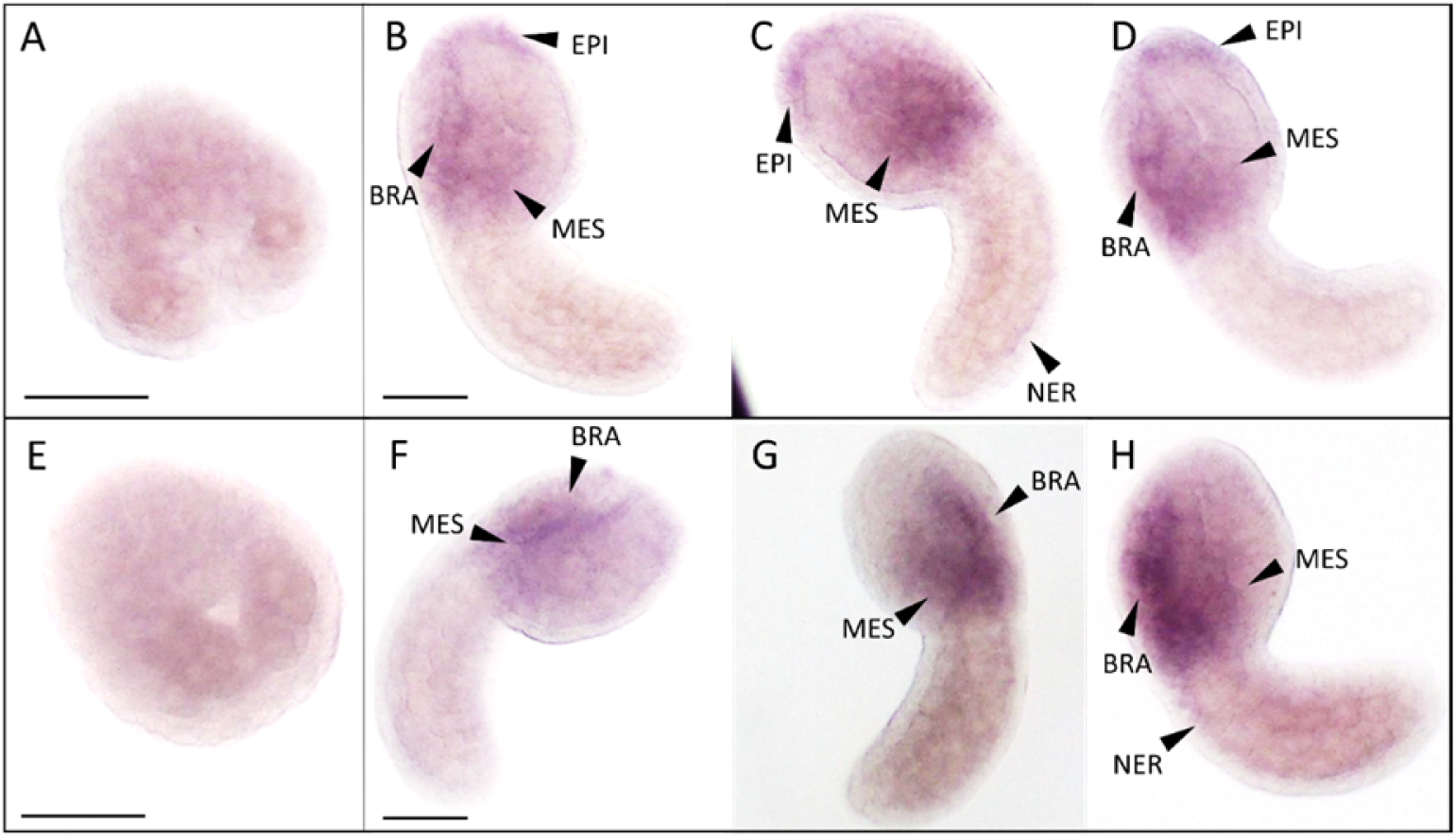
Whole mount ISH on mid gastrula (A, E) and mid tailbud II (B-D, F-H) embryos, with *cr-tiarx1* (A-D) and *cr-g3bp2* (E-H) riboprobes, developed in control condition (A-B, E-F) or in presence of Cd (C, G) or Fe (D, H). EPI: epidermis; MES: mesenchyme; BRA: brain; NER: nerve cord. Scale bar: 100 µm.

### qRT-PCR CONFIRMS A HIGHER mRNA ACCUMULATION BY IRON

Since whole mount ISH results confirmed that *cr-tiarx1* and *cr-g3bp2* were not transcribed at mid gastrula stage, their gene expression, together with that of *cr-ttp*, were quantified in mid tailbud II and hatching larva embryos, both in control condition and after metal exposure (Figure 6). The highest transcription levels were observed after Fe exposure at hatching larva stage for all the studied genes. Except for *cr-tiarx1*, at mid tailbud II stage, a statistically significant increase (p < 0.05) in gene expression in respect to controls was always present.

**Figure 6.**
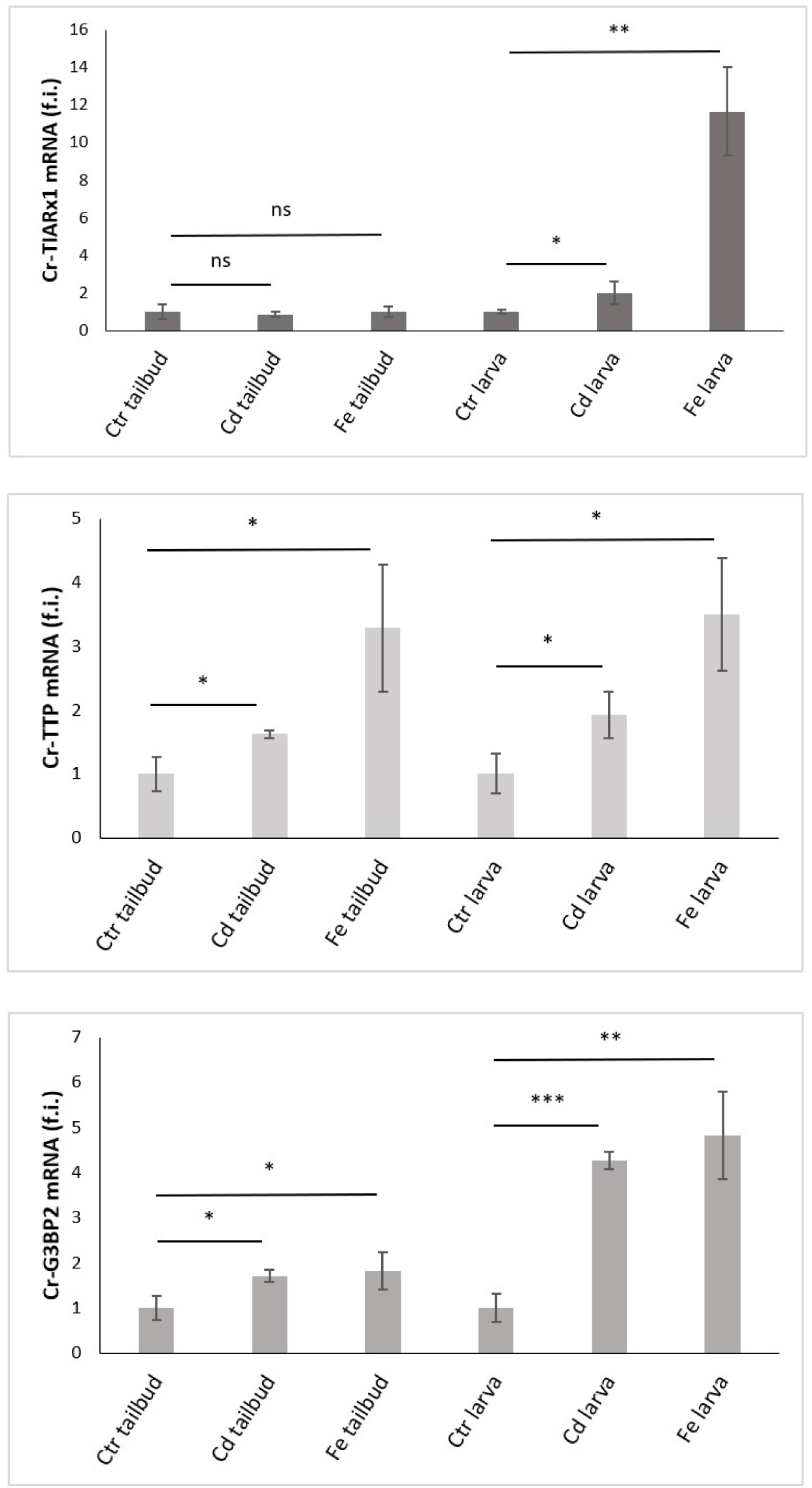
Relative expression levels (fold induction, f.i.) of Cr-TIARx1, Cr-TTP and Cr-G3BP2, in pooled mid tailbud II embryos (tailbud) and hatching larvae (larva), developed in FSW without metal (Ctr) or with cadmium (Cd) or iron (Fe). Asterisks mark significant differences with respect to controls (***: p < 0.001; **: p < 0.01; *: p < 0.05; ns: p > 0.05).

### CRISPR/Cas9 KNOCKOUT OF SG GENES GREATLY AFFECTS MESENDODERMAL DERIVATIVES

Gel images resulting from Genomic Cleavage Detection assay, on genomic DNA extracted from initial tailbud embryos electroporated with Eef1a>Cas9 together with U6>tiar_sgRNA3, U6>ttp_sgRNA4 or U6>g3bp_sgRNA2 vectors, are reported in Figure S3. From the digestion reactions of re-annealed PCR amplicons, we were able to identify parental bands and cleavage products of the expected sizes. SgRNA3, targeting *cr-tiarx1*, caused 23.8% of genetic modification in DNA; the percentage amounts to 14.8% for sgRNA4 targeting *cr-ttp*, and 20.3% for sgRNA2 targeting *cr-g3bp2*. Controls obtained from genomic DNA extracted from embryos electroporated with Eef1a>Cas9 vector alone were also considered: in this case, no cleavage products were detected (data not shown). Gel image conditions were optimized with ImageJ software, to reduce background signal as much as possible. The third cleavage band detected in the case of *cr-tiarx1,* is due to naturally occurring single-nucleotide polymorphisms (SNPs).

Different phenotypes resulting from tissue-specific knockout of *cr*-*tiarx1*, *cr*-*ttp* and *cr*-*g3bp2*, by using the sgRNAs reported above, can be found in Figure 7, 8 and 9 for mid tailbud II stage, and Figure 10 for hatching larva. Control phenotypes, for the two analyzed stages, are reported in Figure S4.

**Figure 7.**
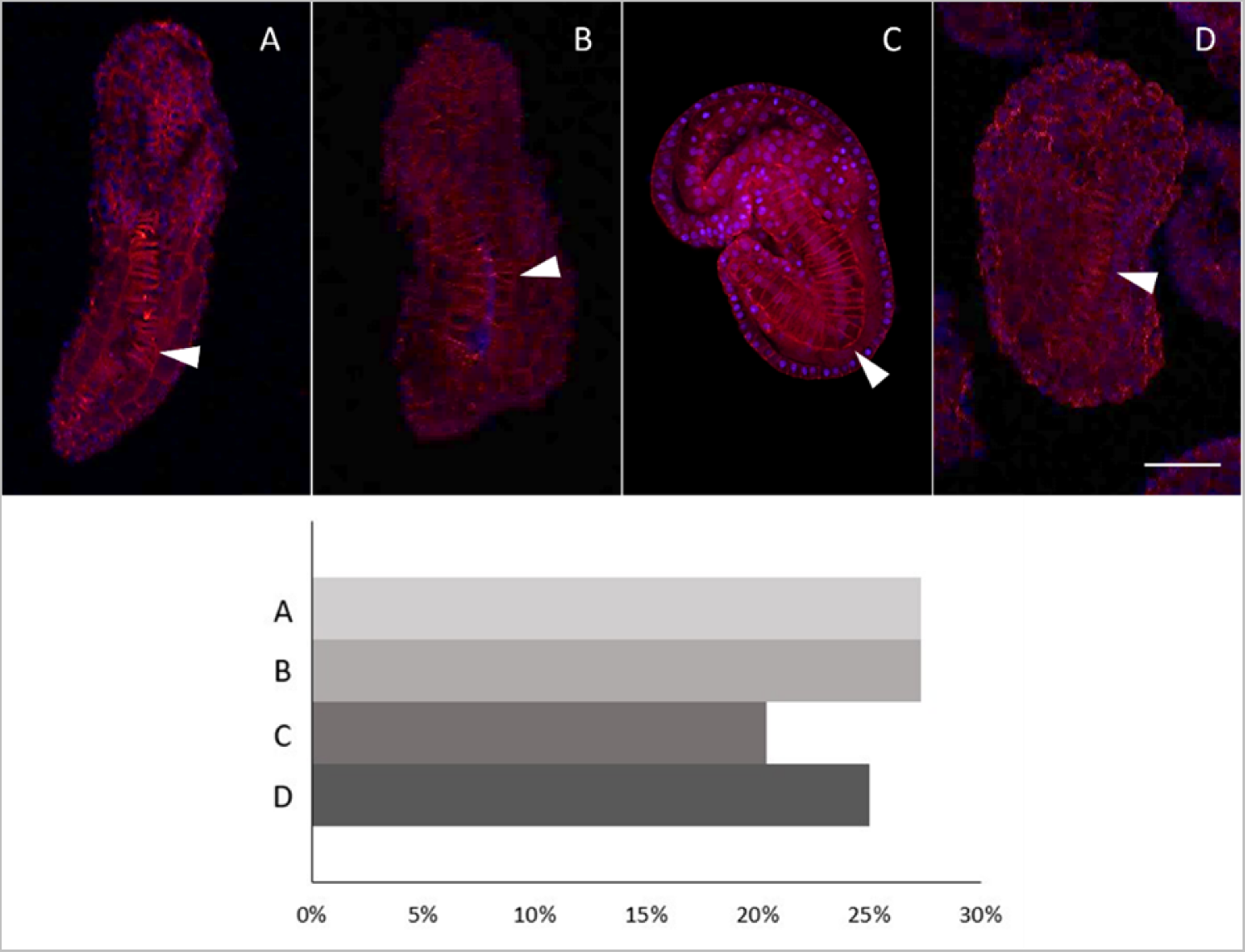
Four different phenotypes (A-D), related to mid tailbud II stage, resulting from *cr-tiarx1* knockout. Arrows indicate the tail and the notochord inside, as seen by Phalloidin staining (red) and nuclei labeled with DAPI/Hoechst (blue). Scale bar: 100 μm. The frequency (%) of phenotype is also reported.

**Figure 8.**
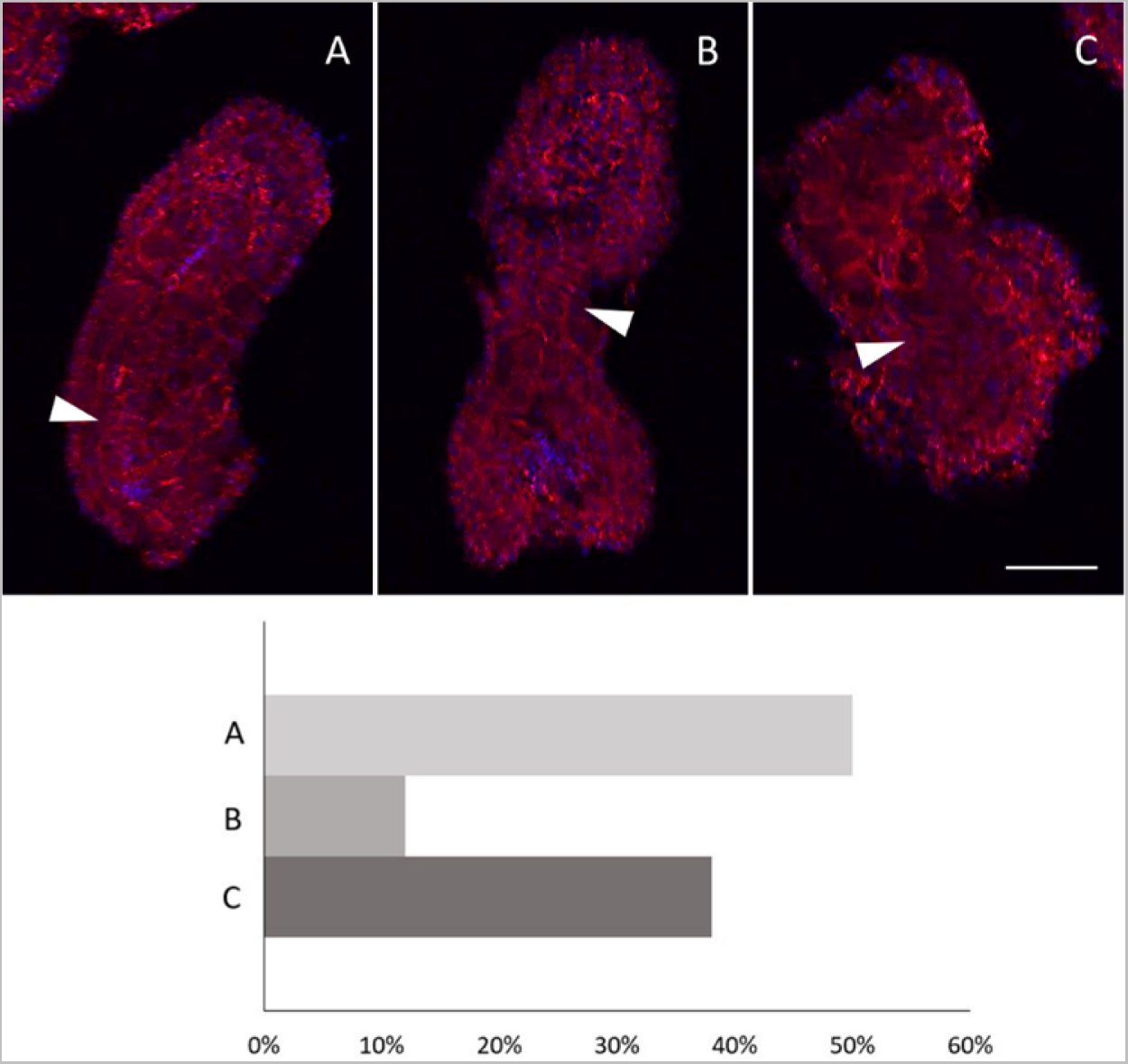
Three different phenotypes (A-C), related to mid tailbud II stage, resulting from *cr-ttp* knockout. Arrows indicate the tail and the notochord inside, as seen by Phalloidin staining (red) and nuclei labeled with DAPI/Hoechst (blue). Scale bar: 100 μm. The frequency (%) of phenotype is also reported.

**Figure 9.**
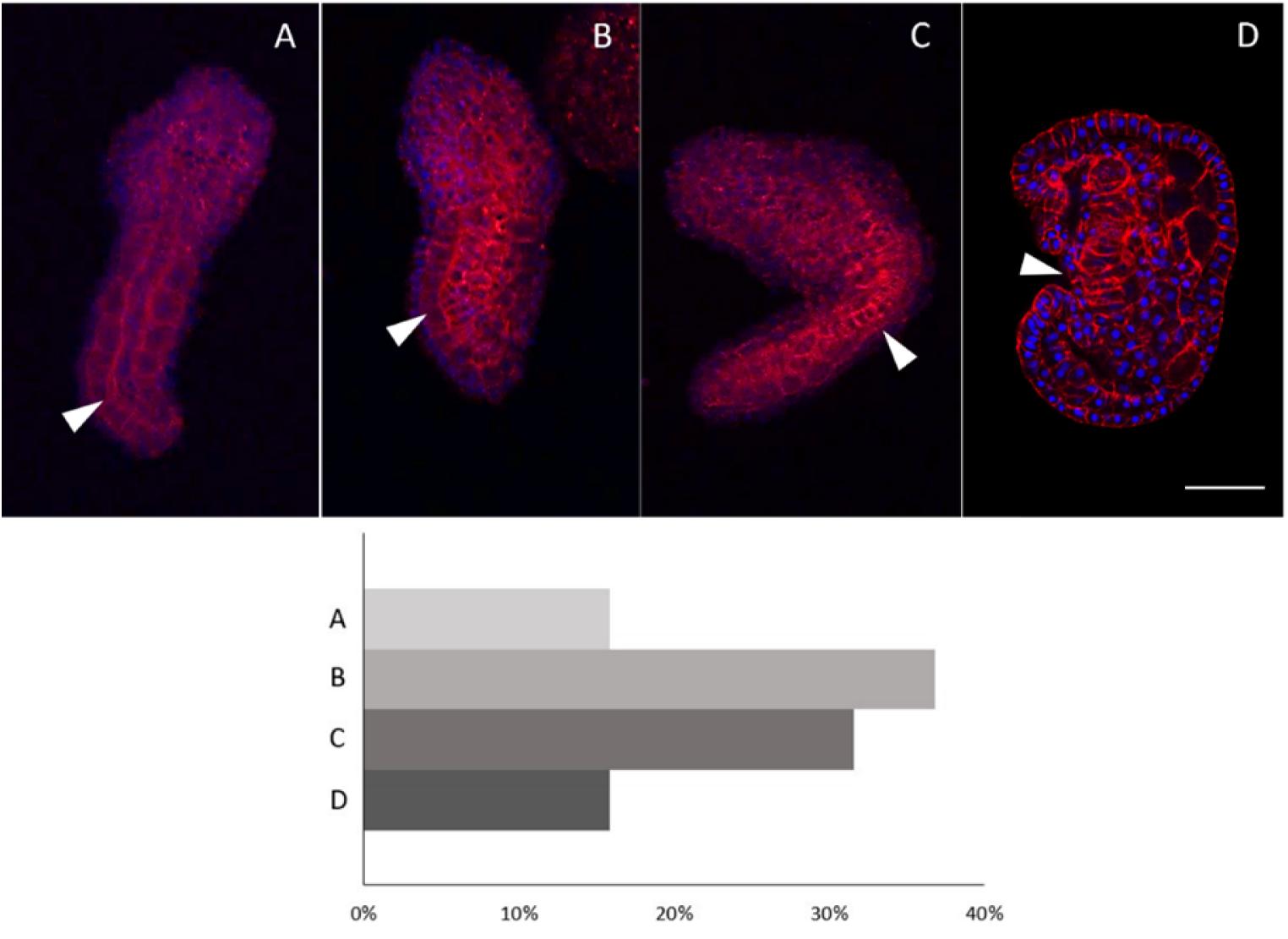
Four different phenotypes (A-D), related to mid tailbud II stage, resulting from *cr-g3bp2* knockout. Arrows indicate the tail and the notochord inside, as seen by Phalloidin staining (red) and nuclei labeled with DAPI/Hoechst (blue). Scale bar: 100 μm. The frequency (%) of phenotype is also reported.

**Figure 10.**
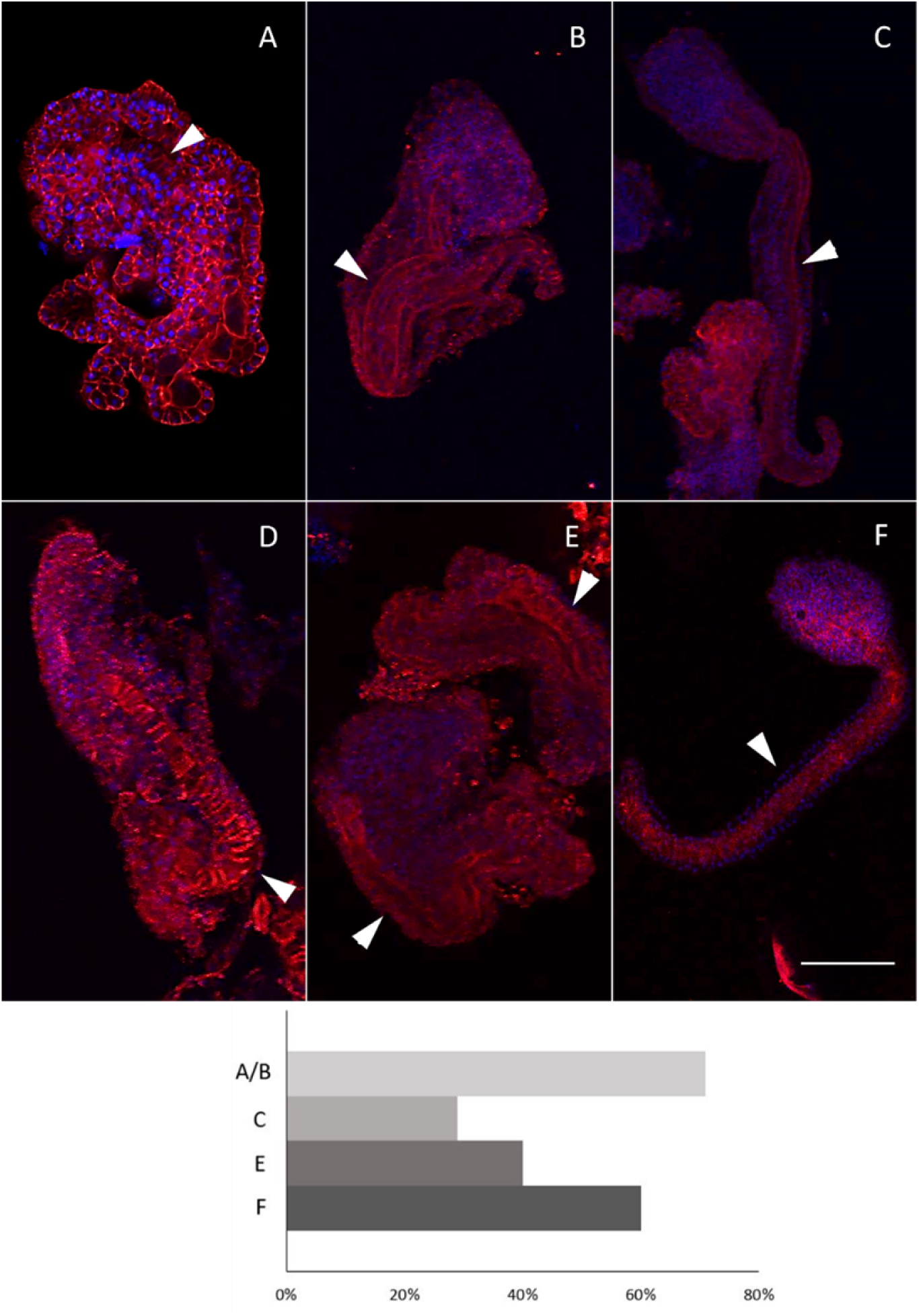
Six different phenotypes (A-F), related to hatching larva stage, resulting from *cr-tiarx1* (A, B, C)*, cr-ttp* (D) or *cr-g3bp2* (E, F) knockout. Arrows indicate the tail and the notochord inside, as seen by Phalloidin staining (red) and nuclei labeled with DAPI/Hoechst (blue). Scale bar: 100 μm. The frequency (%) of phenotype is also reported.

After *cr-tiarx1* knockout, the most abundant phenotypes present in the mid tailbud II embryos were those reported in Figure 7A (27%, n=44) and 7B (27%), with malformations also involving the proximal part of the tail, due to a light curvature of the notochord, which was more pronounced in 20% of embryos (Figure 7C), and an enlargement of cerebral vesicle area. Finally, 25% of embryos showed an uncontrolled cell proliferation, resulting in larger embryos, with the head not well distinguished from the tail (Figure 7D), as already noted in phenotype B in the mid-terminal region of the tail.

The phenotype mostly presents after *cr-ttp* knockout (38%, n= 30) is that reported in Figure 8A, very similar to phenotype B obtained after *cr-tiarx1* knockout (Figure 7B). In 12% of embryos, the uncontrolled cell proliferation caused a bifurcation of the terminal tail (Figure 8B). After *cr-ttp* knockout, 50% of embryos did not look like as a tailbud stage (Figure 8C) but were similar to the phenotype C obtained after *cr-tiarx1* knockout (Figure 7C).

*cr-g3bp2* knockout caused curvature of the tail, in different points, as shown by phenotype A (16%, n= 34), B (37%) and C (32%) (Figure 9). The major malformation was observed in 16% of embryos (Figure 9D), like phenotypes reported in Figure 7C and 8C.

As regards hatching larva stage, specimens with a well-distinguished tail, even if larger than normal, were observed after *cr*-*tiarx1* and *cr*-*g3bp2* knockouts in 29% (n=59) and 60% (n= 40) of the observed larvae, respectively (Figure 10C, F). In all the other cases, the embryos were highly malformed, with uncontrolled proliferation and the tail region was identified by the presence of the notochord (Figure 10A, B, D, E). *cr*-*ttp* knockout resulted in embryos (n= 40) with a single type of phenotype (Figure 10D), showing uncontrolled proliferation and the tail region marked by the presence of the notochord.

## DISCUSSION

The maintenance of homeostasis is essential for ensuring the survival of the organisms and, consequently, the perpetuation of species. This is especially true for organisms living in very variable environmental conditions, such as lagoons, marinas, or harbors. Recent studies have confirmed the ability of ascidians (sessile species of tunicates, the sister group of vertebrates) to cope with stressful conditions through the activation of both antioxidant (Franchi et al., 2012; Ferro et al., 2018) and immune responses (Parrinello et al., 2015; Franchi and Ballarin, 2017; Peronato et al., 2020; Marino et al., 2023). These responses are mainly detectable in hemocytes, representing the detoxification system of ascidians (Franchi et al., 2011, 2014).

SGs are ribonucleoproteic cytoplasmatic foci controlling and modulating the global protein synthesis (Drago et al., 2021; Drago et al., 2023). TIAR, G3BP and TTP are three molecular components of SGs which are deeply studied in vertebrates. The aggregation of TIAR and G3BP is fundamental for SG formation, as it leads to temporary AU-rich mRNA silencing thus to ensure the normal cell proliferation and differentiation. An increasing number of reports indicate that SGs exert their action not only in the presence of stressful conditions but whenever cells require a proper control of protein synthesis. As early development and differentiation need a strict control of cell activity, it is conceivable that SGs play a role in these processes. The role of SGs in embryogenesis is still poorly known although some data, indicating their involvement in the above processes, have been collected from studies in mice (Ramos et al., 2004; Zekri et al., 2005; Sánchez-Jiménez and Izquierdo, 2013; Kharraz et al., 2010; Liu et al., 2023) and in *C. elegans* among invertebrates (Huelgas-Morales et al., 2016).

With the present work we aim to increase the knowledge on the role of the *cr*-*tiarx1*, *cr*-*ttp* and *cr*-*g3bp2*, orthologues of mammalian *tiar*, *ttp* and *g3bp*, in early development of a chordate invertebrate, the solitary ascidian *C. robusta*, in control conditions and after metal-induced stress conditions.

After electroporation of vectors with promoters for *cr-ttp*, *cr-tiarx1* and *cr-g3bp2*, we were able to visualize in which tissue the regulation of the gene expressions takes place. As regards mid tailbud II stage, mesenchyme is the mainly involved tissue, in all the experimental conditions, whereas under metal (Fe and Cd) stress conditions the regulation of the studied gene expressions is activated in cells of the nervous system, and, for *cr-ttp*, also in endoderm and in muscle cells.

Mesenchyme cells, located in the trunk, are fated to become hemolymph cells and body-wall muscle in the adult (Passamaneck and Di Gregorio, 2005). Our previous studies confirmed that the orthologues of *tiarx1* and *ttp* are expressed in circulating immunocytes of the solitary ascidian *C. robusta* (Drago et al., 2021) and the colonial species *B. schlosseri* (Drago et al., 2023). Circulating immunocytes of ascidians control both phagocytosis and inflammation, the two typical cellular responses of innate immunity. We already demonstrated that, in *B. schlosseri*, the microinjection in the colonial circulation of a specific anti-TIAR antibody significantly reduces the capability of phagocytes to ingest foreign target cells and of cytotoxic cells to degranulate and trigger the inflammatory response (Drago et al., 2023). In *C. robusta*, transcripts for TIAR and TTP are present mainly in granulocytes, an immunocyte type that can act as a detoxifying organ, able to accumulate metal ions and to migrate to the tunic, as observed also *in B. schlosseri*, where they undergo apoptosis and clearance by tunic phagocytes (Franchi et al., 2014; Drago et al., 2021, 2023). ISH results show that also *cr-g3bp2* is transcribed in immunocytes, thus strongly suggesting its involvement in immune responses (Figure S5).

Endodermal cells, present in the trunk and in the tail (endodermal strand), will originate the endostyle, the branchial sac and the digestive organs of the adult (Hirano and Nishida, 2000). Our previous studies confirmed *tiarx1* transcription and the expression of the relative protein in the endostyle, stomach and intestine of *B. schlosseri* (Drago et al., 2023). As regards *C. robusta*, *cr-tiarx1* and *cr-ttp* transcription upon metal exposures (Cu, Zn and Cd), were confirmed, by qRT-PCR, in the intestine, which is one of the first the organs contacting xenobiotics upon filter-feeding (Drago et al., 2021). Supplementary results, in Figure S6, confirm that also *cr-g3bp2* is expressed in the intestine of *Ciona* adults, with a maximum of transcriptional activity at 24 and 72 h and a minimum at 48 h of exposure to Cd. This trend was already observed for the other two genes and is likely related to the post-transcriptional control operated by SGs: once perceived the acute stress, at 48 h, cells disassemble their SGs to unlock the translation of mRNAs for anti-stress proteins.

In mid tailbud II stage, the transcription of all the studied genes was observed also in the tail (endodermal strand, nerve cord, notochord, muscle cells), which is not confirmed in later development. Indeed, hatching larvae are very close to metamorphosis during which the tail will be resorbed and will not contribute to the formation of the adult body: this is probably the reason of the absence of transcription of *tiarx1*, *ttp* and *g3bp2*. Conversely, the activation of the studied genes in hatching larval stage is visible in the endoderm, for *cr-g3bp2* and *cr-ttp*, and mesenchyme, for *cr-tiarx1* and *cr-ttp*, especially in embryos exposed to Fe and Cd.

The central nervous system of the ascidian larva includes a visceral ganglion and a sensory vesicle in the trunk, connected by a neck and a caudal nerve cord (Olivo et al., 2021). Our data, indicating the activity of *cr-ttp* promoter in the neural plate of the mid-gastrula embryos, suggest the involvement of TTP in the development of the ascidian brain. This observation agrees with the reported transcriptional activity of G3BP2, TIAR and TTP in the mammalian brain, likely implicated in both signal transduction and RNA-metabolism (Kennedy et al., 2001; Rayman and Kandel, 2017; Li et al., 2019). The activity of the promoter in brain precursors is increased by metal exposure in both the mid-gastrula and mid tailbud II embryos: this suggests a decisive role of these genes in controlling cell homeostasis and facing stressful conditions.

Our whole mount ISH results confirmed that the transcription of the studied genes occurs mainly in brain, mesenchyme, and endoderm of mid tailbud II embryos, remarking the possible role of *cr-tiarx1*, *cr-ttp* and *cr-g3bp2* in adult immune responses and in defense of the adult digestive system in addition to development. We also quantified the transcriptional activity of the three genes in mid tailbud II embryos and hatching larvae by qRT-PCR, in untreated and metal-exposed specimens. Except for *cr-tiarx1* at mid tailbud II stage, every studied gene showed a statistically significant increase in gene transcription in metal exposed samples with respect to the controls in both the considered stages, reaching the highest level after Fe exposure. According to our previous hypothesis, on post-transcriptional control operated by SGs formed by TIARx1, G3BP2 and TTP, an acute stress should lead to a decline in transcription of the relative genes, to allow the unlock of mRNA from disassembled SGs to codify proteins to face the adverse conditions. A possible explanation, of the observed increase in expression during metal-induced stress conditions, is that cells need to codify the key proteins for the assembly of SGs, in which to store AU-rich mRNAs for proteins active in the control of cell proliferation and differentiation and cellular metabolism, rather than proteins active in attenuating stress conditions; AU-rich mRNAs for anti-stress proteins, such as metallothionein and glutathione (Franchi et al., 2011, 2012), could be, conversely, free in the cytoplasm and promptly translated. SGs could act, in stressed embryos, in control of mRNAs for TNF, p53, STAT5B and MYC, which protection is valuable to ensure a normal embryogenesis (Piecyk et al., 2000; Johnson and Blackwell, 2002; Jackson et al., 2006; Marderosian et al., 2006; Liao et al., 2007; Vignudelli et al., 2010; Geng et al., 2015; Takayama et al., 2018; Zhang et al., 2021).

The importance of *cr-tiarx1*, *cr-ttp* and *cr-g3bp2* in controlling cell proliferation and differentiation is highlighted by our CRISPR/Cas9 experiments. The knockout of the above genes, in absence of stress, affected normal embryogenesis, leading to malformations at level of the proximal part of the tail, due to unnatural curvatures or uncontrolled cell proliferation, and the head/trunk, sometimes showing an enlargement of cerebral vesicle area, often undistinguished from each other especially in hatching larvae, in respect to controls. In the case of *cr-g3bp2*, the knockout seemed to cause an arrest in head/trunk development at late tailbud III stage, and this is confirmed by the few studies present in the literature, according to which the inhibition of G3BP causes growth retardation or embryonic lethality, due to an increase in neuronal cell death by apoptosis (Zekri et al., 2005).

## CONCLUSIONS

This represents the first study on SG molecular markers in ascidian embryogenesis. Phenotypes resulted from transfected embryos of *C. robusta* revealed that *cr-tiarx1*, *cr-ttp* and *cr-g3bp2* are required to ensure normal embryogenesis, also ensured by the formation of SGs during stress conditions, to allow the control of mRNA translation in proteins regulating cell proliferation, differentiation and metabolism.

## MATERIALS AND METHODS

### ANIMALS

*C. robusta* is an invertebrate chordate belonging to the subphylum Tunicata, widely used in embryogenetic studies since it allows to obtain hundreds of embryos, after a single artificial fertilization event, which develop up to the larval stage within a day, ready to adhere to a solid substrate and metamorphose into adults (Passamaneck and Di Gregorio, 2005; Hotta et al., 2007; Hotta et al., 2020). In addition, *Ciona* embryos can be easily electroporated with specific constructs for mutagenic experiments (Stolfi et al., 2014; Fujiwara and Cañestro, 2018; Zeller, 2018; Papadogiannis et al., 2022). *Ciona* represents an excellent model organism also for its annotated compact genome, easily accessible in the ANISEED database, with relatively short intergenic regions which facilitate the identification of *cis*-regulatory elements for the control of gene expression (Dardaillon et al., 2020).

### OLIGONUCLEOTIDE DESIGN

Oligonucleotides for gene expression studies (qRT-PCR) and gene knockout experiments (CRISPR/Cas9) were designed in the coding region (CDS) of *tiar* and *ttp* (Drago et al., 2021), and *g3bp* (https://www.aniseed.fr/; transcript ID: KY2019:KY.Chr1.2038.v1.SL1-1). The last sequence was confirmed by sequencing the amplicons after amplification (Eurofins Genomics Europe Shared Services GmbH, Ebersberg, Germany). Intron/exon compositions were then analyzed (http://ghost.zool.kyoto-u.ac.jp/SearchGenomekh.html).

Oligonucleotides for the preparation of single guide RNAs (sgRNAs) were designed using CRISPOR (Concordet and Haeussler, 2018) with a Doench’16 cutoff of ≥ 60. In order to obtain efficient knockout, we placed the sgRNAs in genomic areas covering functional domains. Off-target sequences with less than three mismatches were avoided as well as sequences with four or more consecutive thymine near the 3’ end, as they can cause termination of transcription by RNA polymerase III. Oligos containing the sgRNA sequences were ordered with overhangs complementary to the vector sequence (U6>sgRNA F+E, addgene59986) and to initiate the transcription of sgRNAs at the U6 promoter (Stolfi et al., 2014). Primers for the validation of genomic DNA cleavage, achieved by CRISPR/Cas9 constructs, were designed to flank the target locus so that the potential cleavage site was not in the center of the amplicon.

For gene reporter assay, primers were designed to isolate potential *tiar*, *ttp* and *g3bp cis*-regulatory regions, identified in the ghost database genome browser (http://ghost.zool.kyoto-u.ac.jp/default_ros.html) genome browser in areas of high chromatin accessibility (staged, whole embryo wild type ATAC-seq data) and upstream of the transcription start site. The IDT OligoAnalyzer tool (https://eu.idtdna.com/pages/tools/oligoanalyzer) was used to check all the designed primers, which were synthesized by Merck Life Science S.r.l. (Milano, Italy), Eurofins Genomics Japan (Tokyo), and Microsynth (Balgach, Switzerland) (Table S1).

### G3BP GENE AND PROTEIN ORGANIZATION

Gene structure of *C. robusta g3bp* was analyzed by matching the cDNA, in part obtained by sequencing and then reconstructed *in silico* through ANISEED database, with the genomic sequence (http://ghost.zool.kyoto-u.ac.jp/SearchGenomekh.html). For sequence comparison studies and to check to have correctly named the analyzed sequence of *C. robusta*, we used the tool BLAST of NCBI (https://blast.ncbi.nlm.nih.gov/Blast.cgi). For the study of G3BP protein organization, orthologous amino acid sequences of some metazoans were aligned together with that of *C. robusta*, using the Clustal Omega software (https://www.ebi.ac.uk/Tools/msa/clustalo/). On the multi-alignment, domain architecture, previously predicted with the SMART program (http://smart.embl-heidelberg.de/smart/set_mode.cgi?NORMAL=1), was visualized. The LALIGN tool (https://www.ebi.ac.uk/Tools/psa/lalign/) was used to compare G3BP sequence of *C. robusta* with each orthologous sequence considered in the multi-alignment, to evaluate the degree of identity and similarity. G3BP protein molecular weight was calculated with Expasy Compute pI/Mw tool (https://web.expasy.org/compute_pi/).

### PLASMID PREPARATION

The In-Fusion primer design tool (Takarabio.com) was used to design and simulate the cloning procedures. *tiar* (530 nt upstream the CDS), *ttp* (318 nt upstream the CDS) and *g3bp cis*-regulatory regions, selected with primers containing specific overhangs (Table S1), were introduced in the pSP72-1.27 vector driving LacZ (Corbo et al., 1997), through a ligation reaction performed with the In-Fusion Snap Assembly (Takara Bio, Japan) master mix (50°C for 15 min). Q5 High-Fidelity polymerase (NEB) was used to linearize the backbone vector (LaZ_IF_Fw and LacZ_IF_Rev primers, Table S1) and the potential cis-regulatory regions, starting from genomic DNA extracted from sperms (see paragraph below). The PCR products were gel-purified with Monarch DNA Gel Extraction Kit (NEB) and used for the ligation reactions. Ligation products were cloned in JM109 competent cells (Promega, USA) and screened by colony PCR with OneTaq Quick-Load 2X Master Mix (NEB). Positive colonies were cultured overnight (o/n) for miniprep, performed according to the PureYield Plasmid Miniprep System (Promega) instructions, and confirmed by sequencing (Microsynth, Switzerland). The three constructs were then cultured for 24 h at 37°C and purified with NucleoBond Xtra Midi Kit (Macherey-Nagel, Germany) and used for gene reporter assay.

In the present study, we produced ten sgRNA vectors, targeting *tiar*, *ttp* and *g3bp*, to guide Cas9 nuclease, according to Stolfi et al. (2014). U6>sgRNA(F+E) purified plasmid, linearized with *Bsa*I (New England Biolabs, Japan), was ligated with double-stranded DNA (Table S1) (vector to insert ratio 1:2), annealed at 10 µM (90 ℃ for 5 min, 30℃ for 30 min), using T4 DNA ligase (Promega), o/n at 4°C.

For purification of digested U6>sgRNA(F+E), the gel slice was dissolved in NaI solution (gel slice to NaI ratio 1:3) containing glass powder. Once gel was completely dissolved by mixing, EasyTrap buffer (50% ethanol, 100 mM NaCl, 10 mM Tris-HCl, pH 7.5) and 1 mM EDTA was used to wash the glass powder. Then, DNA was separated from glass powder by heating at 55°C in nuclease free water and centrifugation. Ligation products were cloned in DH-5α *Escherichia coli* chemically competent cells, plasmids were purified from positively screened colonies with QIAprep Miniprep Kit (QIAGEN) and sequenced at the Research Institute for Molecular Genetics of Kochi University. Therefore, the constructs were cloned again in TOP10 competent cells, which were cultured for 24 h, and finally purified with QIAGEN Plasmid Midi Kit and used for CRISPR/Cas9 experiments.

FoxD>Cas9 plasmid was prepared starting from FoxD>LacZ plasmid (Mita and Fujiwara, 2007). The primers to amplify the upstream region of *FoxD*, with Takara Ex Premier DNA Polymerase (Takara Bio), are reported in Table S1. EF1α>Cas9 (addgene59987) (Stolfi et al., 2014) was digested with NheI (NEB) and SnaBI (NEB), to excise out the upstream region of EF1α, then the PCR-amplified *FoxD* upstream region and restriction-digested plasmid vector were gel-purified, by using the EasyTrap buffer as described above, and finally fused by a recombination reaction using In-Fusion HD Cloning Kit (Takara Bio). In-Fusion reaction solution was directly used for transformation of TOP10 strain of *E. coli*.

### GENOMIC DNA EXTRACTION

Genomic DNA was extracted from *Ciona* sperm. SDS lysis buffer (1% SDS, 50 mM NaCl, 10 mM EDTA, 50 mM Tris, pH 8), with 0.02 mg/ml of proteinase XIV, was mixed with the sperm o/n at 50°C. Phenol-chloroform-isoamyl alcohol (25:24:1) was added to remove digested proteins and after centrifugation at 12,000xg for 45 min, 3 M Natrium acetate (pH 5.2) was added to the supernatant together with absolute ethanol (EtOH). After DNA precipitation o/n at -20°C, and centrifugation at 12,000xg for 20 min, DNA pellet was washed with 80% EtOH and centrifugated twice, at 12,000xg for 20 min. DNA pellet was air-dried and resuspended in nuclease free water. DNA concentration and purity were assessed with the Nanodrop ND-1000 spectrophotometer (ThermoFisher Scientific).

### EMBRYO COLLECTION AND TREATMENT

Adult specimens of *Ciona robusta* used in this study were collected in Chioggia, in the southern area of the Lagoon of Venice, Italy, and in the Usa area of Tosa city, Kōchi Prefecture, Japan; in the last case, *Ciona* juveniles, kindly provided by Yutaka Satou Lab of Kyoto University, were reared in nature until sexual maturation. Animals were maintained in aerated aquaria, filled with filtered sea water (FSW) or Super Marine Art SF1 artificial sea water (ASW; Tomita Pharmaceuticals), at 18°C under direct light and fed with Phyto Marine (Oceanlife, Bologna, Italy) and Reefpearls (DVH Aquatic, Holland).

Good quality eggs and sperm, from dissected adults, were used for *in vitro* fertilizations (Kari et al., 2016). Sperm was activated with a solution 0.05 M Tris (pH 9.5) in FSW or ASW and added to the eggs. After 15 min, embryos were gently collected, dechorionated with dechorionation solution (1% sodium thioglycolate, 0.05% Proteinase XIV) and 0.05 M NaOH in FSW or ASW and washed in FSW or ASW. About 200 embryos were reared in 10 cm, agarose coated Petri dishes, in order to prevent the adhesion of the embryos and cultured at 18°C in FSW or ASW, with or without (controls) metal ions. Samples were collected at the desired stage, and subsequently used for whole mount ISH and qRT-PCR. For metal treatments, iron (Fe) was used as essential metal, cadmium (Cd) as non-essential metal; after several tests to determine the LC_50_ (lethal dose, 50%), 1 μM of metal ions was chosen as sub-lethal concentration. Storage solutions were previously prepared by dissolving the metal salt (FeCl_2_ or CdCl_2_) in distilled water and the metal concentration was verified, with a Perkin Elmer 4000 Atomic Absorption Spectrophotometer; working solutions were prepared by diluting the storage solutions in FSW.

In another series of experiments (gene reporter and CRISPR/Cas9), dechorionated embryos were immediately subjected to electroporation.

### ELECTROPORATION

Dechorionated embryos were electroporated with a single pulse at 50 V for 16 msec, in a 4 mm electroporation cuvette, containing 0.77 M mannitol solution (Kari et al., 2016). For gene reporter assay we used 40 μg of plasmid, containing *tiar* (pTiar>LacZ), *ttp* (pTtp>LacZ) or *g3bp* (pG3bp>LacZ) cis-regulatory sequences, driving LacZ, and 30 μg of Fog>H2B:mCherry plasmid, marking the successful uptake of the electroporation mix, in a total volume of 350µl (Papadogiannis et al., 2022). For CRISPR/Cas9 experiments we used 75 μg of sgRNA-expressing plasmids for *tiar*, *ttp* or *g3bp* knockout (U6>tiar_sgRNA, U6>ttp_sgRNA, U6>g3bp_sgRNA) and 25 μg of Eef1a>Cas9 or FoxD>Cas9 plasmids. Electroporated plasmid amounts were per 700 μl of total volume.

For gene reporter assay, electroporated embryos were reared in FSW with or without metal ions, as reported above, until the desire stage; for CRISPR/Cas9 experiments, electroporated embryos were maintained in ASW, until the desire stage. Eef1a>Cas9 was used to evaluate CRISPR/Cas9-targeted mutagenesis of *tiar*, *ttp* and *g3bp* locus at initial tailbud stage, considering that the ubiquitous Eef1a promoter is not active before the 64 cell stage (Stolfi et al., 2014), through the GeneArt Genomic Cleavage Detection Kit. Once verified which sgRNA function in gene knockout, FoxD>Cas9 was used in electroporation for tissue-specific (mesendodermal derivatives, including posterior brain and central nervous system, head endoderm, notochord, and trunk ventral cells) mutagenesis of *Ciona* embryos (Imai et al., 2009; Abitua et al., 2012; Gandhi et al., 2017).

### GENE REPORTER ASSAY

Gene reporter assay was carried out to analyze the differential tissue activity of cis-regulatory elements in embryos developed in the presence of metals. Samples at the studied stages (mid gastrula, mid tailbud II and hatching larva) were incubated for 1 h in blocking solution, i.e. 0.25% Tergitol, 5% heat inactivated normal goat serum in phosphate-buffered saline (PBS: 1.37 M NaCl, 0.03 M KCl, 0.015 M KH_2_PO_4_, 0.065 M Na_2_HPO_4_, pH 7.2), and incubated with Anti-Beta-Galactosidase Purified Monoclonal Antibody (Promega), 1:200 in blocking solution for 2 h. After several wash with 0,25% Tergitol in PBS, embryos were incubated in Alexa Fluor 488 goat anti-mouse IgG (H+L) (Invitrogen), 1:500 in blocking solution for 1 h. Embryos were washed and mounted in Vectashield (Vector Laboratories) and observed under a Leica TCS SP5 Confocal microscope.

### CRISPR/Cas9 KNOCKOUT: DNA CLEAVAGE DETECTION ASSAY AND PHENOTYPE STUDY

The DNA cleavage validation for *tiar*, *ttp* and *g3bp* was assessed with GeneArt Genomic Cleavage Detection Kit (Thermofisher), starting from genomic DNA extracted from initial tailbud embryos co-electroporated with specific sgRNAs and Eef1a>Cas9 before the first cell division. Following cleavage, indels were created in genomic DNA by normal cellular repair mechanisms, so loci, where the double-strand breaks were expected, were PCR-amplified (primers in Table S1). PCR products were denatured and reannealed and mismatches were subsequently detected and cleaved by the kit Detection Enzyme. Negative controls were also considered, for each sgRNAs, by adding nuclease free water instead of the detection enzyme. Resultant bands were visualized by electrophoresis and analyzed with ImageJ software. The absence of cleavage was also verified in embryos electroporated with Eef1a>Cas9 only.

For each gene, at least one sgRNA vector was identified as active in the DNA cleavage and used in the CRISPR/Cas9 experiments co-electroporated with FoxD>Cas9, to mediate tissue-specific mutagenesis and evaluate the importance of TIAR, TTP and G3BP for a correct embryo development. Electroporated embryos, fixed at the desired stage (mid tailbud II and hatching larva), in 2% paraformaldehyde (PFA) in ASW with 0.1% Tween-20 for 15 min, were blocked in 1% powdered milk in PBS containing 0.1% Tween-20 (PBST) for 30 min. Embryos were treated with 100 nM Rhodamine Phalloidin (Thermo Fisher Scientific) or Acti-stain 488 fluorescent Phalloidin (Cytoskeleton, Inc.) in PBST for 20 min, in dark conditions, followed by 1 µg/ml DAPI (Thermo Fisher Scientific) or Hoechst (Life Technologies corporation) in PBST for 15 min. Embryos were finally mounted with 86% glycerol and observed under a Leica TCS SP5 or a Nikon C1si confocal microscope. Embryos electroporated with FoxD>Cas9 only were considered as control of the normal phenotype development in the absence of DNA cleavage.

### WHOLE MOUNT ISH

Specific DNA templates were obtained from cDNA libraries (VES103-I09 for TIAR, VES72-L12 for G3BP, VES65-B17 for TTP) (Roure et al., 2007). Kanamycin-resistant colonies were selected and cultured o/n purified with Plasmid Miniprep system (Promega) and verified by sequencing (Microsynth, Switzerland). PCR products with flanking T7 promoters were used as template for the synthesis of anti-sense riboprobes using a T7 RNA polymerase (Roche) (Mercurio et al. 2023). Riboprobes were purified by precipitation with 0.1 M LiCl and 75% EtOH. After centrifugation at maximum speed for 15 min, the pellet was washed with 70% EtOH, airdried and dissolved in nuclease free water.

To verify the correlation between *tiar*, *ttp* and *g3bp* gene expression and gene reporter assay results, whole mount ISH was carried out at the mid gastrula and mid tailbud II stages as previously described (Mercurio et al. 2021). Briefly, control and metal treated embryos, were fixed (4% PFA, 0.5 M NaCl, 0.1 M MOPS) for 1 h, dehydrated with EtOH and stored in 70% EtOH at -20°C until use. After rehydration in EtOH, embryos were permeabilized with 2 µg/ml proteinase K in PBST at 37°C for 5 min, fixed again for 1 h and pre-hybridized in hybridization solution, i.e. 50% formamide, sodium citrate solution 5x (SSC 20x: 3 M NaCl, 0.34 M sodium citrate, pH 4.5), 1 µg/ml yeast tRNA, 0.25 µg/ml heparin, and 0.1% Tween-20 in nuclease free water, at 52°C for 2 h. Hybridization was carried out at 52°C for 18 h. After several washes (50% formamide, SSC 5x, 0.1% Tween-20 in nuclease free water) at 52°C, embryos were incubated in 25% normal goat serum in PBST for 2 h. Incubation with anti-digoxigenin-AP Fab antibody (Roche), 1:2000 in PBST, was performed o/n at 4°C. After several washes in PBST, embryos were washed with APT buffer (0.1 M Tris-HCl, pH 9.5, 0.15 M NaCl, 0.1% Tween-20 in nuclease free water) and with APTMg buffer (0.1 M Tris-HCl, pH 9.5, 0.15 M NaCl, 0.05 M MgCl_2_, 0.1% Tween-20 in nuclease free water) for 10 min. Finally, the samples were incubated in NBT (nitro-tetrazolium blue)-BCIP (5 bromo-4-chloro-3’-indolyphosphate p-toluidine) (Sigma-Aldrich), 0.033% in APTMg in dark condition. Embryos were fixed for 1 h and mounted in 86% glycerol for microscope observation. Images were captured with a Leica DM6 B Fully Automated Upright Microscope System.

### TOTAL RNA EXTRACTION, cDNA SYNTHESIS, AMPLIFICATIONS, AND SEQUENCING

To quantify the transcription of *tiar*, *ttp* and *g3bp*, total RNA was isolated from untreated and metal-treated embryos, at mid tailbud II and hatching larva stages, with the RNA NucleoSpin RNA XS (Macherey–Nagel, Düren, Germany) kit. RNA concentration and purity were assessed with the Nanodrop ND-1000 spectrophotometer (ThermoFisher Scientific) and its integrity was verified through electrophoresis. Reverse transcription from 1 μg of total RNA was performed with the ImPromII (Promega) kit. The obtained cDNAs were used for qualitative and quantitative PCR, respectively according to PCRBIO Classic Taq (PCRBIOSYSTEMS, London, UK) and HOT FIREPol EvaGreen qPCR Mix Plus (ROX) (Solis BioDyne) kit instructions. The melting profile was analyzed, by qRT-PCR, to verify the absence of genomic contamination. For each treatment three pool of embryos from three different artificial fertilization events (biological triplicate), at the desired stages, were analyzed, by running each sample three times (technical triplicate).

Total RNA was extracted also from *Ciona* adult intestine and Cr-g3bp_PCRFw and Cr-g3bp_PCRRv primers (Table S1) were used to amplify the retrotranscribed cDNA. Amplicons were separated by electrophoresis and the corresponding bands were purified with the Wizard SV Gel and PCR Clean-Up System (Promega) kit, ligated in pGEM-T Easy Vector (Promega, Madison, WI, USA), and cloned in DH-5α *E. coli* cells. Plasmid DNA was extracted from positively screened colonies with UltraPrep (AHN Biotechnologie GmbH) kit and sequenced by Eurofins Genomics to verify the *g3bp* transcription in *C. robusta* (for *tiar* and *ttp* see Drago et al., 2021).

### DATA COLLECTION AND STATISTICAL ANALYSES

Relative values were obtained from qRT-PCR using the 2^−ΔΔCt^ Pfaffl (Pfaffl, 2001) mathematical model; the levels of transcription were normalized with respect to the housekeeping gene (*β-actin*; primer in Table S1), to compensate for variations in the amounts of cDNA, and with respect to control (untreated) embryos, for each considered embryo stage and metal. Data were expressed as mean of three biological samples (n = 3) ± standard deviation and statistically compared with the Duncan’s test (Snedecor and Cochran, 1980).

In gene reporter assay, only completely electroporated embryos were considered for confocal microscopy analyses. The percentages of abnormal embryos ± standard deviations were compared with the χ^2^ test.

For validation of the DNA cleavage achieved by CRISPR/Cas9 sgRNAs, the relative proportion of DNA contained in each amplicon of the final gel, was determined with ImageJ software, starting from gel pictures. The cleavage efficiency was calculated as follow, according to kit instructions: Cleavage efficiency = 1– [(1–fraction of cleaved bands) ½]; Fraction of cleaved bands = sum of cleaved band intensities/ (sum of the cleaved and parental band intensities).

## ACKNOWLEDGMENTS

Authors wish to thank members of the Yutaka Satou Lab of Kyoto University for providing *Ciona* juveniles, with the support of University National Bio-Resource Project (NBRP) of the Ministry of Education, Culture, Sports, Science and Technology (MEXT) Japan. This research was supported by the Department of Biology, University of Padova (DOR to LB), Horizon 2020 MSCA (COFUND-DP ARDRE 847681 to UR) and JSPS KAKENHI (Grant Number JP23K05793).

## Abbreviations used in the text

ARE: AU-rich element
ASW: artificial sea water
BCIP: 5 bromo-4-chloro-3’-indolyphosphate p-toluidine
CDS: coding region
CRISPR: clustered regularly interspaced short palindromic repeats
eIF2α: eucaryotic initiation factor 2 alphas
EtOH: ethanol
FSW: filtered sea water
G3BP: ras-GTPase-activating protein SH3-domain-binding protein
GAP: GTPase-activating protein
ISH: *in situ* hybridization
LC50: lethal dose, 50%
NBT: nitro-tetrazolium blue
NTF2: N-terminal nuclear transport factor 2
o/n: overnight
p-bodies: processing bodies
PBS: phosphate-buffered saline
PBST: PBS with Tween-20
PKR: protein kinase
qRT-PCR: quantitative real-time PCR
RNP: ribonucleoprotein
RRM: RNA-Recognition Motif
SG: stress granules
sgRNA: single guide RNA
SNP: single-nucleotide polymorphism
SSC: sodium citrate solution
TIAR: TIA 1 related nucleolysin
TPA: tetradecanoyl phorbol acetate
TTP: tristetraprolin
UTR: untranslated region
Znf: zinc finger

---

**Figure S1.**
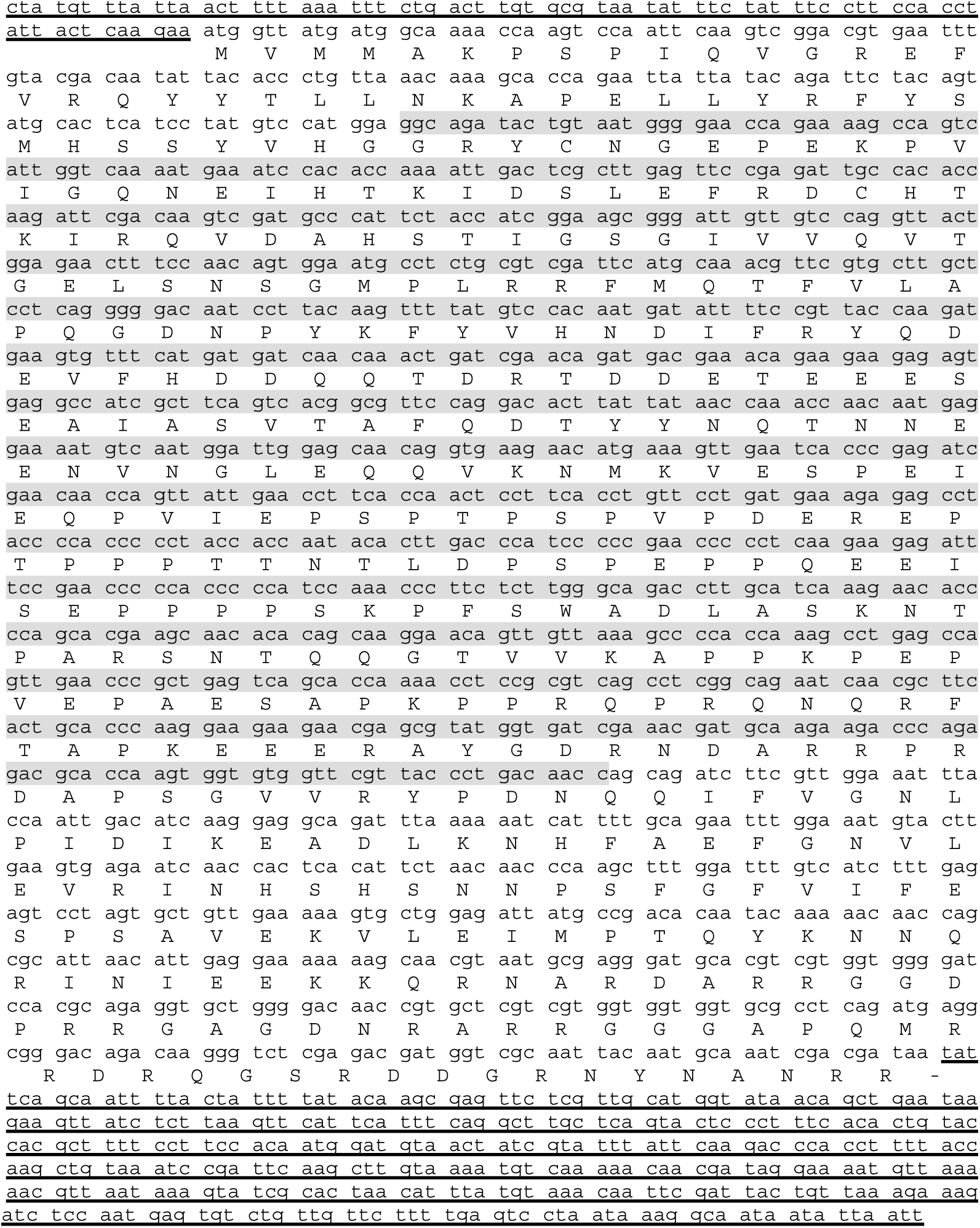
cDNA sequence of Cr-G3BP2 and deduced amino acid sequence. 5’- and 3’-UTR regions are underlined. In grey the sequence obtained with amplicon sequencing.

**Figure S2.**
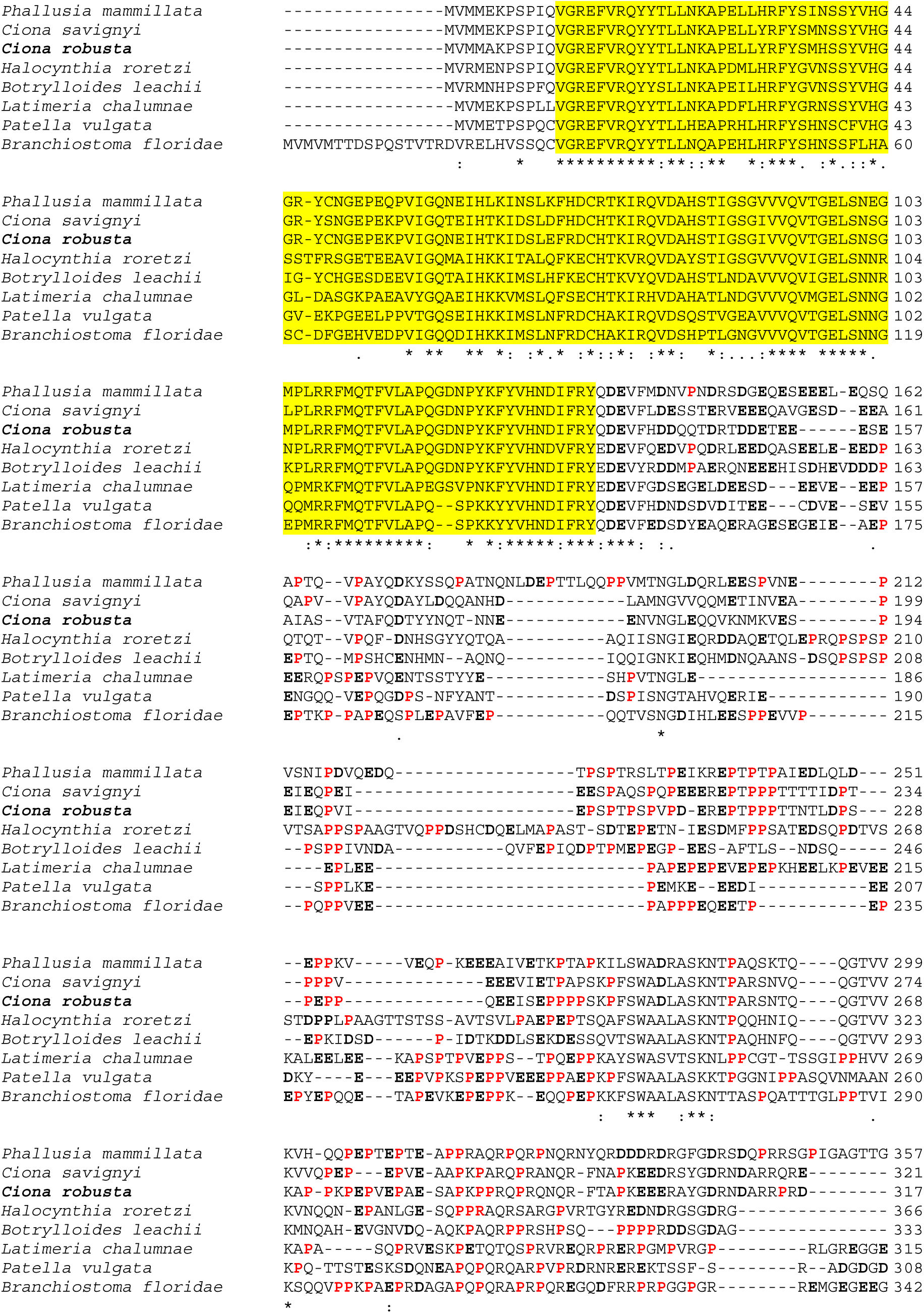
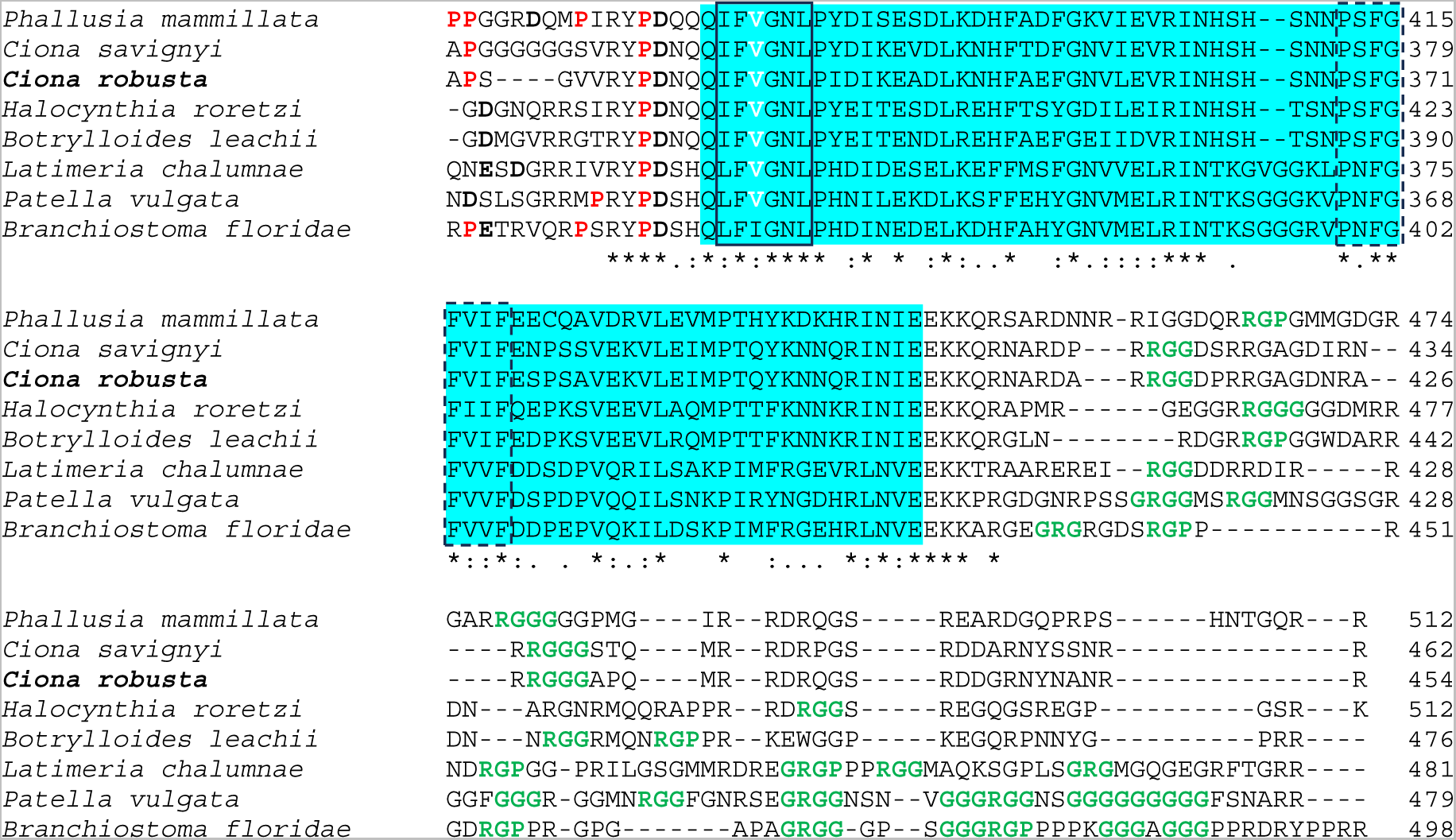
Schematic G3BP2 domain organization. The RRM domain, with RNP2 and RNP1 motifs boxed, respectively, with continuous and a dotted line, is highlighted in blue, whereas the NTF2 domain is highlighted in yellow. In RNP2 motif, the valine residue (V) which allows to discriminate G3BP2 from G3BP1 is in white and bold. The central acid- and proline-rich region shows the glutamic acid (E) and aspartic acid (D) residues, defining the acid-rich region, in black and bold, whereas the proline (P) residues, defining the proline-rich, in red and bold. Closely spaced arginine-glycine repeats (RGG, RGP, GRG, GGG), in C-terminal region, are in green and bold. *= identical amino acids (completely preserved), := very similar amino acids (semi-conservative substitution), .= similar amino acids (conservative substitution); numerals refer to character counts.

**Figure S3.**
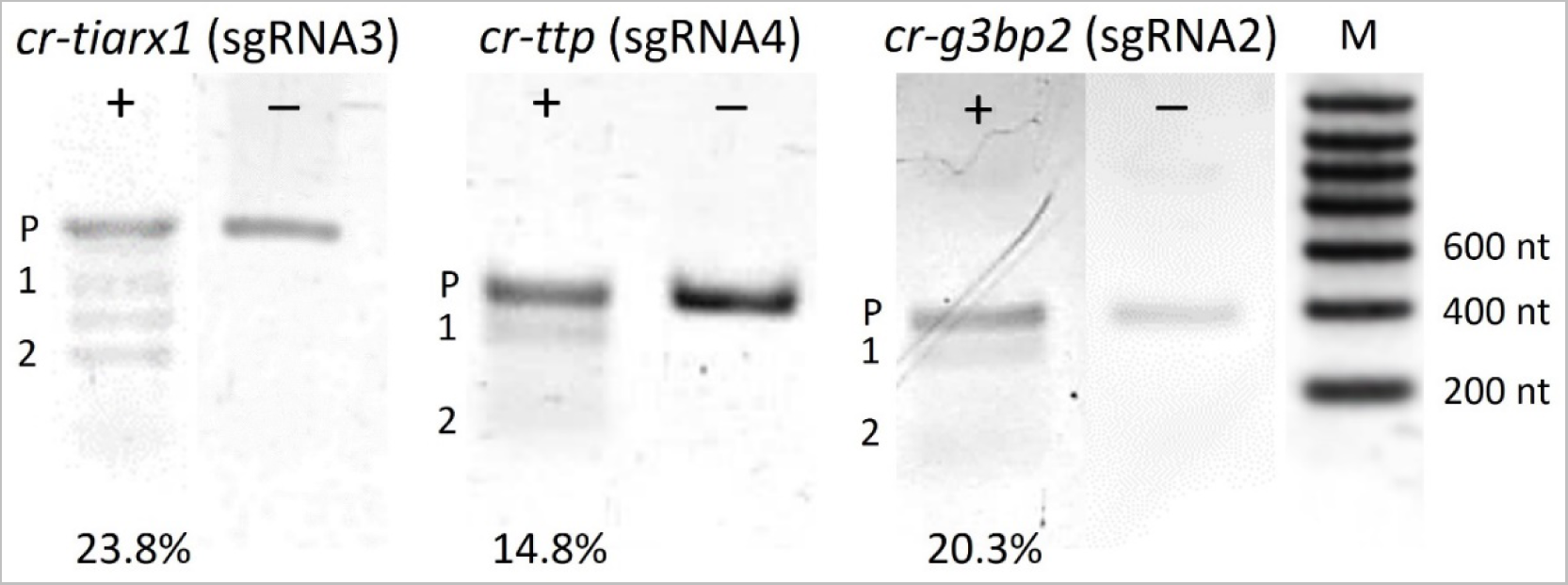
Genomic Cleavage Detection Assay gel analyzed with ImageJ software. Lane profiles for *cr-tiarx1, cr-ttp* and *cr-g3bp2*, referring respectively to sgRNAs 3, 4 and 2, are reported. In lanes “+” are visible the cleavage products, in lanes “-“ are visible the negative controls (no detection enzyme used from the kit). P: parental band; 1: first cleavage product; 2: second cleavage product; M= molecular weight. Cleavage efficiencies (%) are also reported.

**Figure S4.**
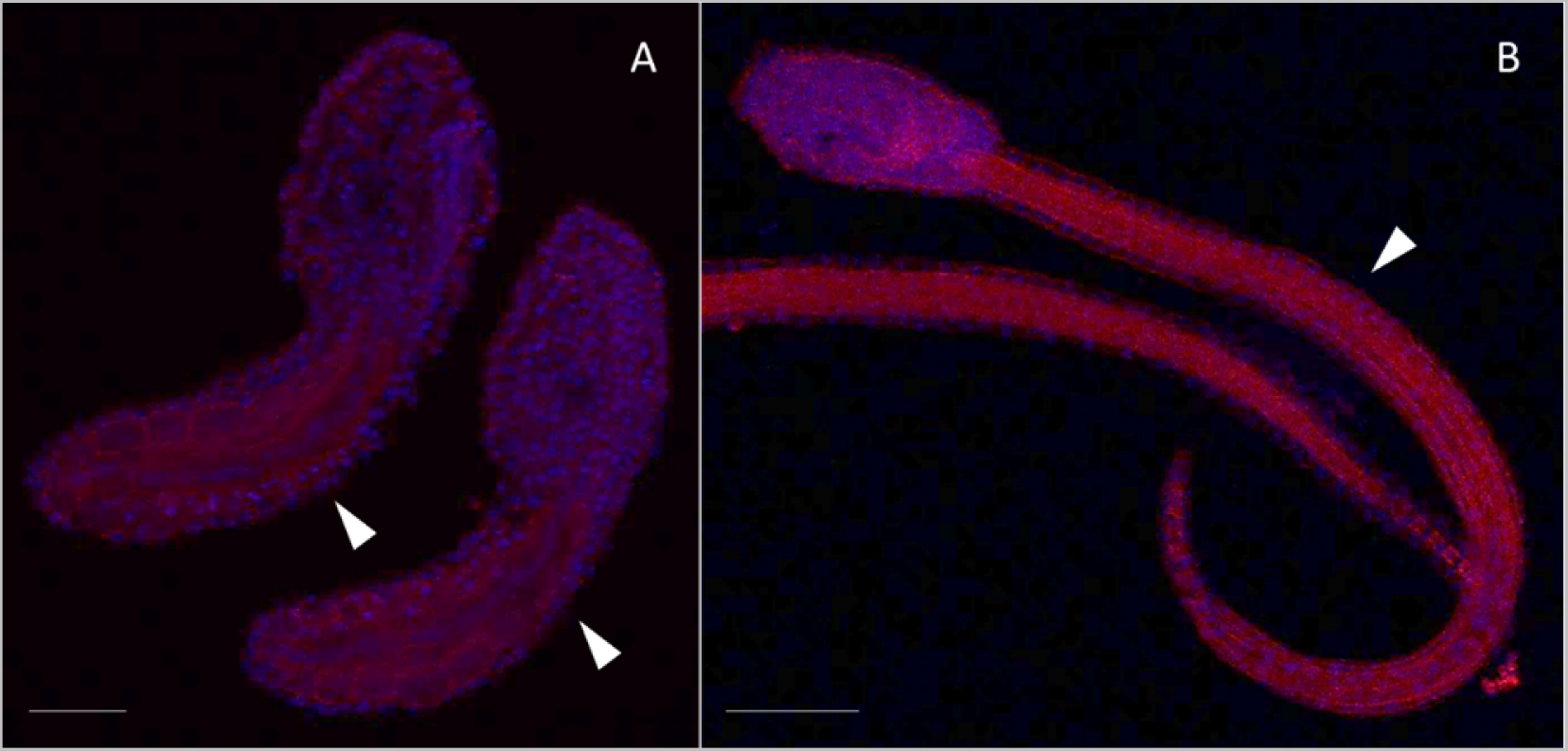
Control phenotypes resulting from electroporation of FoxD>Cas9 only, for mid tailbud II (A) and hatching larva (B) stages. Arrows indicate the tail and the notochord inside, as seen by Phalloidin staining (red) and nuclei labeled with DAPI/Hoechst (blue). Scale bar: 100 μm.

**Figure S5.**
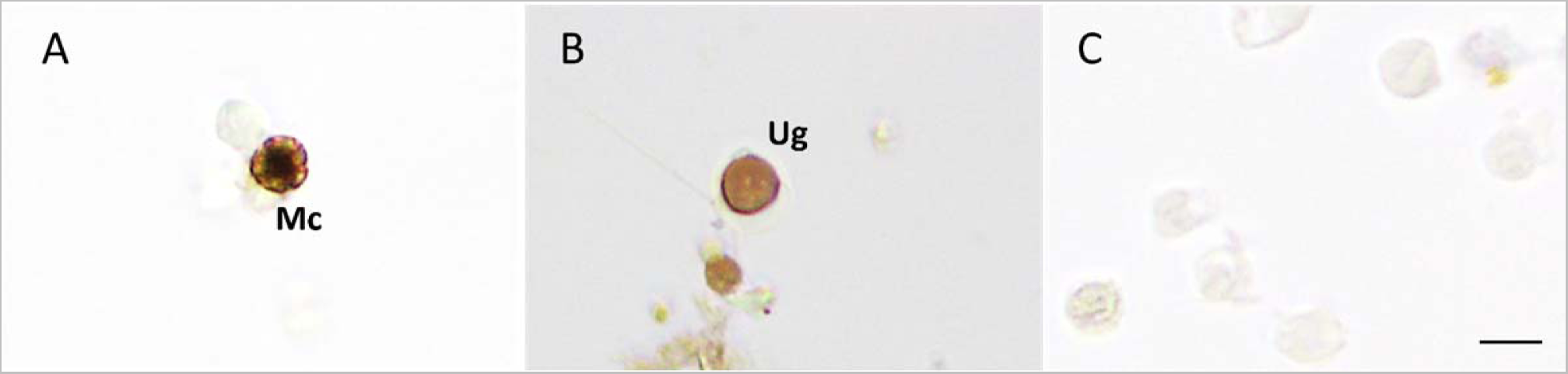
ISH with anti-sense (A, B) and sense (C) *Cr-g3bp2* biotinylated riboprobe, on hemocytes collected from 72 h Cd (10 µM)-treated adults. Mc: morula cell; Ug: unilocular granulocyte. Scale bar: 10 μm.

**Figure S6.**
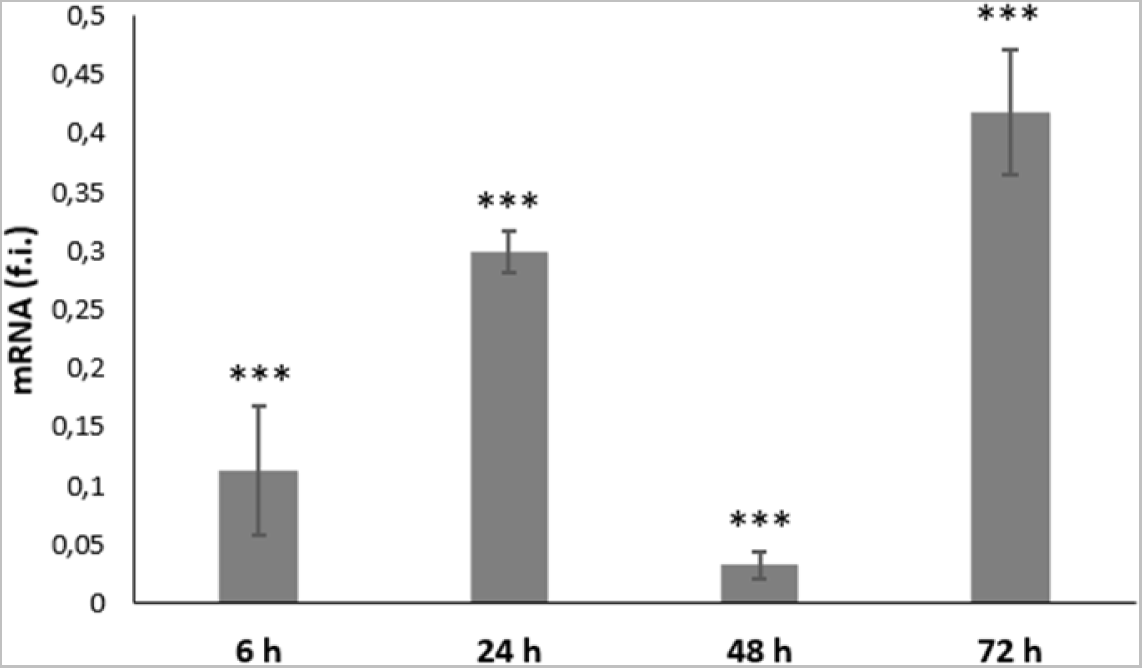
Relative transcription levels (fold induction, f.i.) of Cr-G3BP2, in intestine from adults (n= 3) treated with Cd (10 µM). Data are normalized with respect to controls. Asterisks mark significant differences with respect to controls (***: p < 0.001).

**Table S1.**
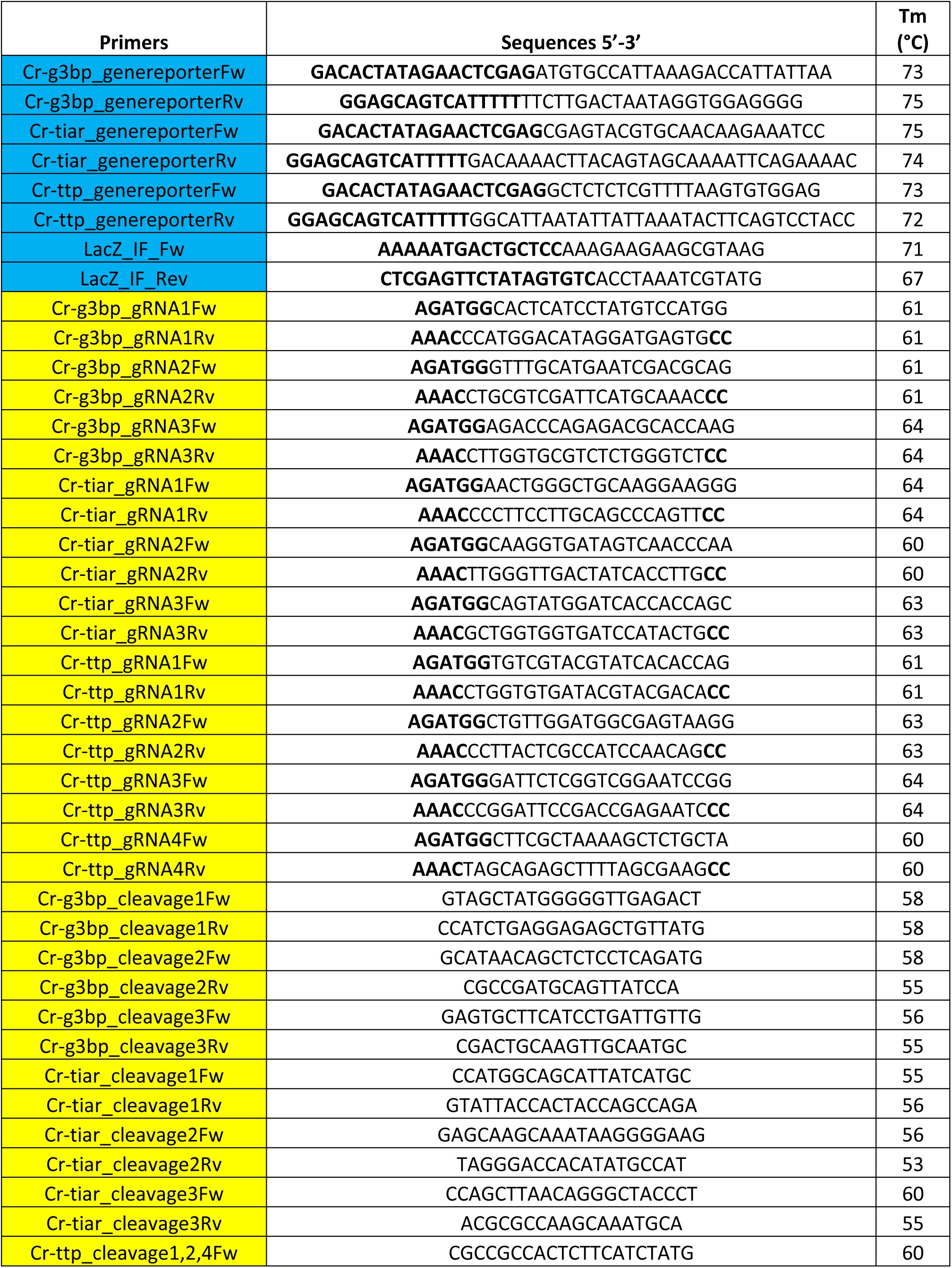
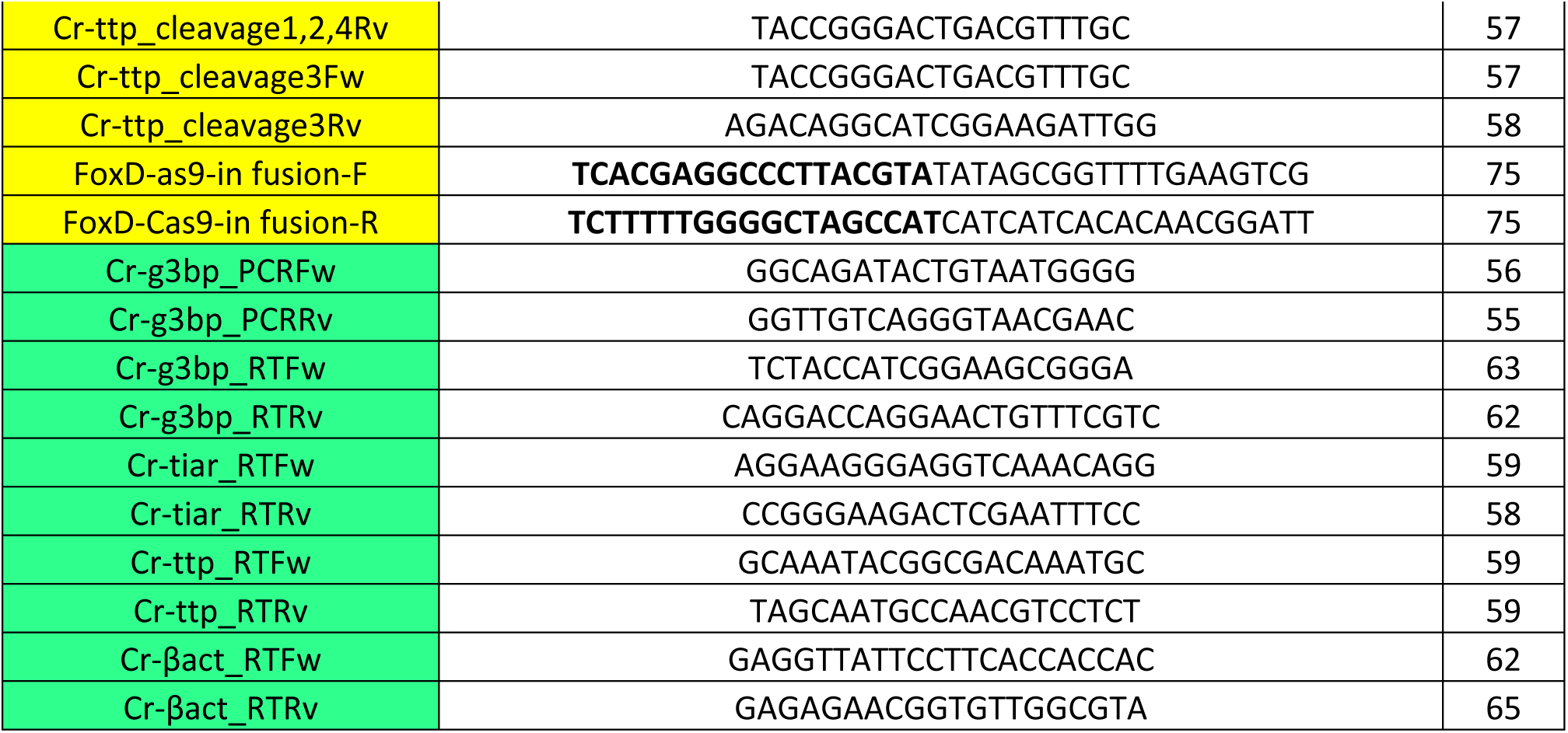
Oligonucleotide sequences, and relative melting temperatures (Tm), used in gene reporter assay (highlighted in blue), CRISPR/Cas9 (highlighted in yellow) and qRT-PCR (highlighted in green). Bold residues represent vector overlapping sequences.

**Table S2.** Percentages of identity and similarity obtained by comparing Cr-G3BP2 amino acid sequence with orthologous sequences of metazoans. E-values, as well as GeneBank (*Branchiostoma floridae, Latimeria chalumnae, Patella vulgate*) and ANISEED accession numbers, are also reported.

| Species | Accession numbers | % identity | % similarity | E-values |
| --- | --- | --- | --- | --- |
| *Ciona savignyi* | Cisavi.CG.ENS81.R27.315248-322198.14078.t | 78.4 | 90.1 | 2.9e-107 |
| *Phallusia mammillata* | Harore.CG.MTP2014.S200.g14562.02.t | 61.0 | 76.7 | 4.9e-76 |
| *Halocynthia roretzi* | Harore.CG.MTP2014.S200.g14562.02.t | 49.2 | 66.9 | 2e-58 |
| *Botrylloides leachii* | Boleac.CG.SB_v3.S147.g02876.01.t | 48.9 | 69.8 | 1.4e-59 |
| *Branchiostoma floridae* | XP_035668730.1 | 47.4 | 68.0 | 1.8e-40 |
| *Latimeria chalumnae* | XP_006010102.1 | 51.6 | 73.1 | 1.3e-49 |
| *Patella vulgata* | XP_050394320.1 | 46.4 | 67.5 | 2.3e-63 |

